# Functional characterization of cardiac fibroblasts in response to hemodynamic alteration

**DOI:** 10.1101/2022.11.05.515315

**Authors:** Manabu Shiraishi, Ken Suzuki, Atsushi Yamaguchi

## Abstract

Excess deposition of extracellular matrix in the myocardium is a predictor of reduced left ventricular function. Although reducing the hemodynamic load is known to improve myocardial fibrosis, the mechanisms underlying reversal of the fibrosis have not been elucidated. We modeled normal myocardium, fibrotic myocardium and myocardium with reduced fibrosis *in vitro*. Fibroblasts differentiated into activated or fibrinolytic types in response to the pericellular environment. Comprehensive gene expression analysis of fibroblasts in each *in vitro* condition showed *Selenbp1* to be one of the genes responsible for regulating differentiation of fibroblasts. *In vitro* knockdown of *Selenbp1* enhanced fibroblast activation and inhibited conversion to the fibrinolytic form. *In vivo* knockdown of *Selenbp1* resulted in structural changes in the left ventricle associated with progressive tissue fibrosis and left ventricular diastolic failure. Selenbp1 is involved in regulating fibroblast differentiation and appears to be one of the major molecules regulating collagen turnover in cardiac fibrosis.

---

Heart failure is the most common reason for hospitalization and a leading cause of death for persons aged 65 years or more, with a mortality rate of approximately 50% within 5 years of diagnosis^1–4^. Left ventricular pressure overload, a companion of systemic hypertension and aortic stenosis, causes progressive interstitial and perivascular fibrosis in association with markedly decreased myocardial compliance, portending a lethal outcome in patients with heart failure with normal systolic function, such as heart failure with preserved ejection fraction^5–8^. Thus, drugs targeting myocardial fibrosis are desirable for treating patients with heart failure.

Normal ventricular function requires preservation of the extracellular matrix (ECM), which functions as a myocardial scaffold, and myocardial fibrosis is characterized by excessive ECM accumulation with activation of cardiac fibroblasts^9, 10^. ECM homeostasis requires continuous synthesis and degradation of matrix proteins (i.e., collagen turnover), a process regulated primarily by fibroblasts resident in the heart^9, 11, 12^. A detailed understanding of the role of fibroblasts in myocardial collagen turnover and of the regulatory mechanisms is of critical importance for establishing cellular and molecular targets for drugs aimed at controlling myocardial fibrosis.

Whereas extensive displacement fibrosis is irreversible, stromal and perivascular fibrosis may be reversible^13^. In a mouse model of ischemic interstitial fibrosis induced by repeated brief episodes of ischemia followed by reperfusion, the fibrosis reversed when ischemic injury was discontinued^14^. In addition, experiments in a rat model suggested that treatment with angiotensin-converting enzyme inhibitor reduces hypertensive cardiac fibrosis^15^. Thus, tissue fibrosis is to some extent reversible by treatment of the underlying disease process.

A myofibroblast is an atypical fibroblast whose precise definition remains debated. Broadly, myofibroblasts can be thought of as activated fibroblasts responsible for the formation of fibrous tissue that leads eventually to heart failure, distinct from fibroblasts that play a role in tissue homeostasis through biomechanical cues (ECM stiffness) and biochemical mediators (such as TGFβ). Myofibroblasts are phenotypically altered fibroblasts that express alpha-smooth muscle actin (αSMA) in stress fibers and accumulate at injury sites. Importantly, although the myofibroblast was initially believed to be the final phase of fibroblast differentiation, we now know that myofibroblasts can deactivate and return to the inactive state characteristic of fibroblasts in homeostatic tissues and that myofibroblasts can differentiate into fibrinolytic cells in the later stages of tissue repair, playing a role distinct from that of progenitor cells^16–18^. Activation of proteases is necessary to remove collagen and other matrix proteins from fibrotic myocardium. Herein, we use the term fibrinolytic cells strictly in reference to cells with increased expression of both matrix metalloproteinase (MMP) 2 and MMP9. The pathways by which myofibroblasts are deactivated and the mechanisms by which myofibroblasts transform into fibrinolytic cells in the heart have not been clarified.

Impeding the development of new targeted therapies is the poorly understood cellular and molecular basis of myocardial fibrosis. This lack of understanding is often attributed to a lack of pathological models. A model that systematically characterizes and recapitulates the specific pathophysiology of myocardial fibrosis is essential. Elasticity of the ECM regulates cell proliferation, apoptosis, and differentiation in various tissues. In particular, stiffened matrices have been shown to promote pathogenic myofibroblast activation^19^. In the study described herein, we created a three-part *in vitro* model that primarily recapitulates human fibrotic myocardium following hemodynamic loading using a medium with an elastic modulus similar to that of myocardial fibrosis, and we investigated phenotypic and functional changes in fibroblasts to explore the underlying regulatory mechanisms. In the process, we searched for key molecules that regulate myocardial collagen turnover, and we validated our findings *in vitro* using RNA interference. We then evaluated the function of identified molecules in an *in vivo* experiment in which heart failure was induced by transverse aortic constriction (TAC).

## Results

### Alterations in the pericellular environment promote differentiation of cardiac fibroblasts into myofibroblasts or fibrinolytic cells

To elucidate the mechanisms by which fibroblast differentiation is regulated, we created a three-part *in vitro* model that recapitulates normal myocardium, fibrotic myocardium following hemodynamic loading (myocardium following HL), and myocardium with reduced fibrosis following hemodynamic loading reduction (myocardium following HLR). Prior to establishing the model, we first tested the pericellular environment (Supplementary Fig. 1a–c) and in so-doing successfully replicated normal myocardium (low elasticity [Low*E*] to Low*E* or Low*E*→Low*E* condition), myocardium following HL (high elasticity [High*E*]+TGFβ to High*E*+TGFβ or High*E*+TGFβ→High*E*+TGFβ condition), and myocardium following HLR (High*E*+TGFβ to Low*E* or High*E*+TGFβ→Low*E* condition) (Fig. 1a). Comparative analysis was then performed. In myocardium following HL, fibroblast activation-related gene (*αSMA*) expression increased about 5-fold, and fibrosis-related gene (*Col1a1*, *Col3a1*) expression increased about 2- and 5-fold, respectively, whereas in myocardium following HLR, expression of *αSMA*, *Col1a1,* and *Col3a1* decreased to levels similar to those in normal myocardium (Fig. 1b). Expression levels of these genes were consistent with protein levels observed by means of quantified immunostaining assay (Fig. 1c). Interestingly, expression of *MMP2* and *MMP9*, which are involved in cardiac ECM degradation, was significantly decreased about 2-fold in myocardium following HL compared to expression in normal myocardium. However, expression of these genes was significantly upregulated in myocardium following HLR compared to that in the other two conditions (Fig. 1d), and this finding was supported by quantified immunostaining (Fig. 1e). In summary, differentiation of fibroblasts to activated form was observed in myocardium following HL, as indicated by increased expression of *αSMA*. However, differentiation to fibrinolytic form was observed in myocardium following HLR, as indicated by increased expression of *MMP2* and *MMP9*, as was deactivation, as indicated by decreased expression of *αSMA*. These results suggest that fibroblasts can change their phenotype by sensing changes in their pericellular environment and that when activated fibroblasts (myofibroblasts) are exposed to normal myocardial conditions, they deactivate and simultaneously acquire a fibrinolytic phenotype.

**Fig. 1.**
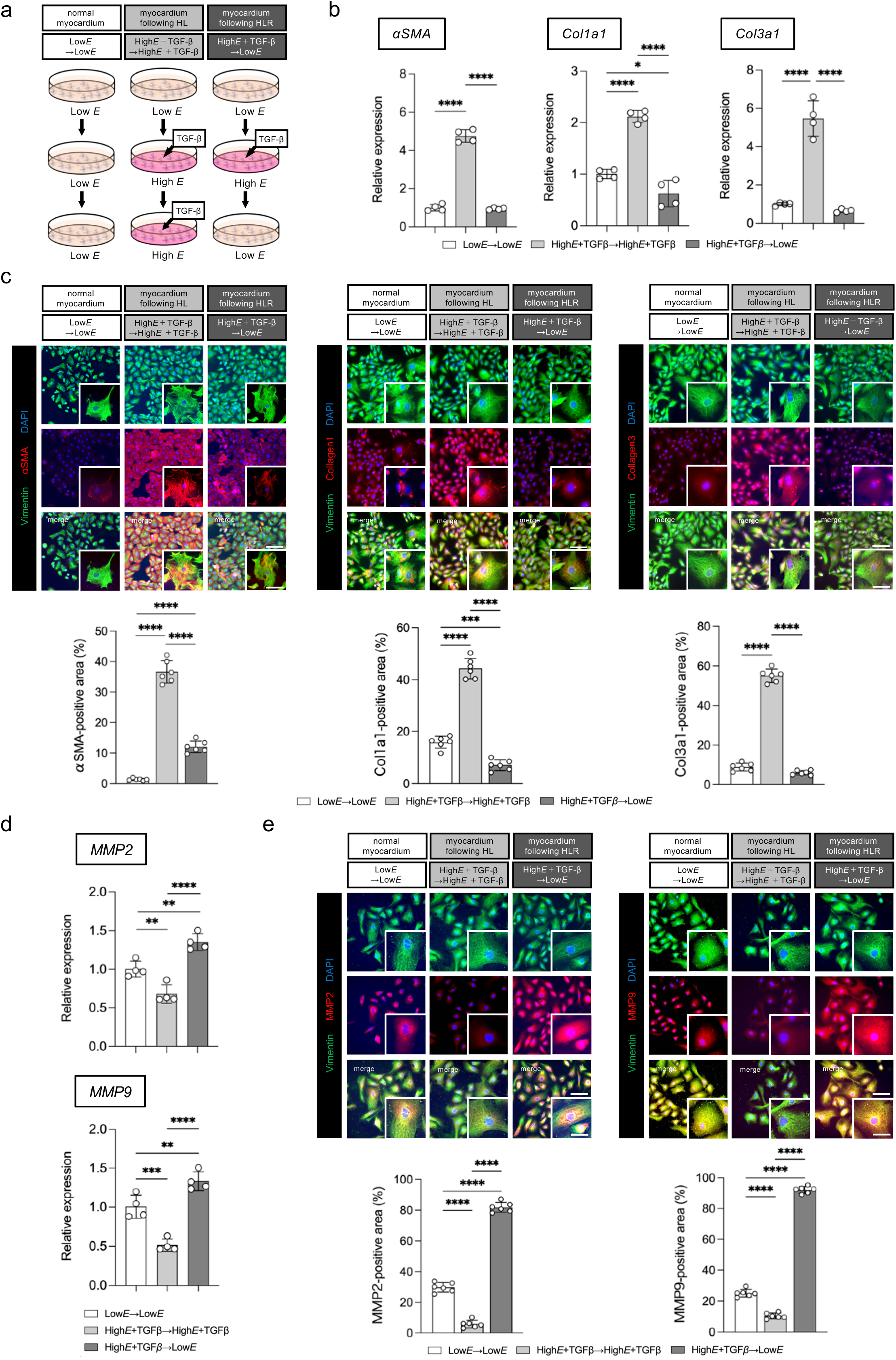
Pericellular environment alters the phenotype of cultured cardiac fibroblasts. *a*, Schematic representation of our three-part *in vitro* model: normal myocardium (Low*E*→Low*E* substrate), myocardium following HL (fibrotic myocardium; High*E*+TGFβ→High*E*+TGFβ substrate), and myocardium following HLR (myocardium with reduced fibrosis; High*E*+TGFβ→Low*E* substrate). *b*, qRT-PCR revealed significant upregulation of fibrosis-associated genes (*αSMA, Col1a1,* and *Col3a1*) in myocardium following HL and downregulation in myocardium following HLR. *c,* Representative images of cardiac fibroblasts stained immunocytochemically for vimentin and αSMA. Activation of cardiac fibroblasts (ratio of vimentin^+^ and αSMA^+^ myofibroblasts to vimentin^+^ fibroblasts) was markedly increased in myocardium following HL. This increase in activation was somewhat attenuated in myocardium following HLR. *d*, qRT-PCR revealed significant downregulation of genes associated with dissolution of fibrosis (*MMP2* and *MMP9*) in myocardium following HL but upregulation in myocardium following HLR. *e*, Representative images of cardiac fibroblasts stained immunocytochemically for MMP2 and MMP9. Expression of MMP2 and MMP9 was markedly decreased in myocardium following HL but markedly increased in myocardium following HLR. *b*, *d*, *n=*4 biologically independent samples per condition; *c*, *e*, Scale bars: 50 μm (low magnification), 10 μm (high magnification), *n=*6 biologically independent samples per culture condition; mean±s.e.m. values are shown. \**P*<0.05, \*\**P*<0.01, \*\*\**P*<0.005, \*\*\*\**P*<0.001 versus the other culture condition(s) by one-way ANOVA.

### Cardiac fibroblasts are induced to differentiate into myofibroblasts and fibrinolytic cells via suppression and promotion of the apoptotic signaling pathway

Comprehensive gene expression analysis of cultured fibroblasts was performed to elucidate the molecular mechanisms involved in the regulation of fibroblast differentiation associated with changes in the pericellular environment. Biologically relevant sets were configured to compare gene expression in fibroblasts in myocardium following HL against that in fibroblasts in normal myocardium (Set A) and gene expression in fibroblasts in myocardium following HLR against that in fibroblasts in myocardium following HL (Set B). Genes whose expression increased more than 1.5-fold numbered 960/24,834 (3.87%) and 512/24,834 (2.06%) in Set A and Set B, respectively. Genes whose expression decreased more than 1.5-fold numbered 837/24,834 (3.37%) and 512/24,834 (1.55%), respectively. The search for known pathways associated with fibroblast differentiation revealed that the TGFβ signaling pathway and the MAPK signaling pathway were significantly activated in Set A, whereas the oxidative stress response, apoptotic signaling pathway, senescence, and autophagy were significantly suppressed (Fig. 2a). In Set B, where myofibroblasts showed deactivation and differentiation into fibrinolytic cells, the TNF-alpha signaling pathway and apoptotic signaling pathway were significantly activated, whereas the TGFβ signaling pathway was significantly suppressed (Fig. 2a). The apoptotic signaling pathway was suppressed during fibroblast activation (Set A) but promoted during fibroblast deactivation and fibrinolytic differentiation (Set B). Assuming that fibroblast activation and differentiation into the fibrinolytic form are regulated by activation or inhibition of a common pathway, we found change to the apoptotic signaling pathway common to both Set A and Set B, and thus the apoptotic signaling pathway was deemed a candidate pathway. In addition, supportive of the apoptotic signaling pathway as the candidate pathway for regulating functional change in fibroblasts, enrichment analysis of the genes with significantly altered expression in Set A and Set B also revealed GO:0010942: positive regulation of cell death as a molecular function among the top 30 most variable items (Supplementary Fig. 2a, b). Furthermore, functional classification of the genes with variable expression showed apoptosis-related clusters to be relatively large in both Set A and Set B (Supplementary Fig. 2c, d; Supplementary Table 1). To evaluate the progression of fibroblast apoptosis with respect to the three conditions, we compared the enzymatic activity of caspase in fibroblasts. Activity of caspase-3/7,-6, and -9 was significantly decreased in myocardium following HL but significantly increased in myocardium following HLR (Fig. 2b), a finding that was confirmed by cleaved-caspase-3 fluorescent immunocytochemical staining (Fig. 2c). As described above, fibroblast apoptosis was suppressed with the change from normal myocardium to myocardium following HL but promoted with the change from myocardium following HL to myocardium following HLR. To confirm that suppression and promotion of the apoptotic signaling pathway are part of the regulatory mechanism underlying phenotypic change in fibroblasts, caspase-3 inhibitor Z-DEVD-FMK was added to fibroblasts under each of the three conditions (Supplementary Fig. 3a). Addition of Z-DEVD-FMK to normal myocardium suppressed caspase-3 activity in fibroblasts (Fig. 3d). Addition of Z-DEVD-FMK to myocardium following HL and to myocardium following HLR significantly increased expression of fibrosis-related genes (*αSMA, Col1a1, Col3a1*), and these patterns were supported by results of quantified immunocytochemistry (Fig. 2e, Supplementary Fig. 3b). However, expression of fibrinolysis-related genes (*MMP2 and MMP9*) was significantly decreased by the addition of Z-DEVD-FMK, especially to myocardium following HLR, as assessed by quantitative reverse transcription-polymerase chain reaction (qRT-PCR) and quantified immunocytochemistry (Fig. 2f, Supplementary Fig. 3c). Caspase-3 inhibition in cultured fibroblasts promoted fibroblast activation in myocardium following HL. However, following HLR, caspase-3 inhibition prevented deactivation of myofibroblasts while attenuating their differentiation into fibrinolytic cells. These results suggested association between the apoptotic signaling pathway and the regulation of fibroblast differentiation in response to changes in the pericellular environment.

**Fig. 2.**
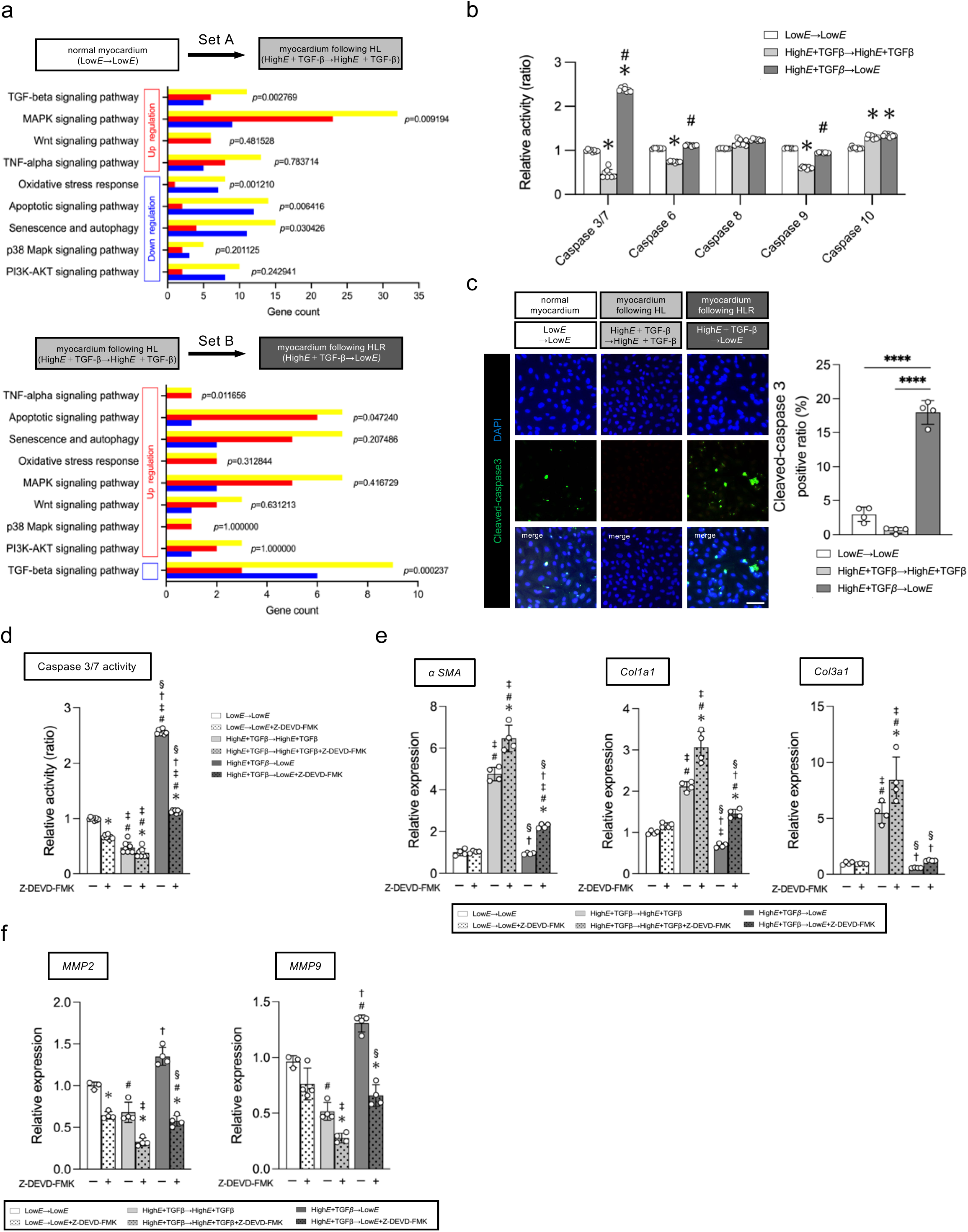
Activation and inhibition of the pathway that regulates apoptosis of cultured cardiac fibroblasts affect their differentiation into myofibroblasts and fibrinolytic cells. *a*, (Upper panel) Results of comprehensive microarray analysis of gene expression of fibroblasts cultured on a substrate mimicking the elastic modulus of myocardium following HL (fibrotic myocardium; High*E*+TGFβ→High*E*+TGFβ substrate) versus that of fibroblasts cultured on a substrate mimicking the elastic modulus of normal myocardium (Low*E*→Low*E* substrate) (Set A). (Lower panel) Results of comprehensive microarray analysis of gene expression of fibroblasts cultured on a substrate mimicking the elastic modulus of myocardium following HLR (myocardium with reduced fibrosis; High*E*+TGFβ→Low*E* substrate) versus that of fibroblasts cultured on a substrate mimicking the elastic modulus of myocardium following HL (fibrotic myocardium; High*E*+TGFβ→High*E*+TGFβ substrate) (Set B). Pathways associated with fibroblast activation are shown. *P* values are shown for the variation in each signaling pathway. Pathways with significant changes in gene expression are indicated by yellow bars. Of the genes with significant changes in expression, those with elevated expression are indicated by red bars, and those with decreased expression are indicated by blue bars. *b*, Enzyme activity of caspase-3/7, -6, -8, -9, and -10 was significantly decreased in fibroblasts of myocardium following HL compared to that in fibroblasts of normal myocardium. Caspase-3/7 activity was significantly increased in fibroblasts of myocardium following HLR, whereas caspase-6 and -9 activity was comparable to that in fibroblasts of normal myocardium. *c,* Representative images of cardiac fibroblasts stained immunocytochemically for cleaved-caspase-3. Expression of cleaved-caspase-3 was markedly increased in myocardium following HLR. *d,* Caspase-3/7 enzyme activity was significantly decreased in the High*E*+TGFβ→High*E*+TGFβ condition and increased in the High*E*+TGFβ→Low*E* condition. Z-DEVD-FMK significantly decreased caspase-3/7 enzyme activity in all conditions. *e,* qRT-PCR revealed that Z-DEVD-FMK significantly upregulated fibrosis-associated genes (*αSMA, Col1a1,* and *Col3a1*) of fibroblasts in myocardium following HL (fibrotic myocardium; High*E*+TGFβ→High*E*+TGFβ substrate). *f,* qRT-PCR revealed that Z-DEVD-FMK significantly downregulated genes associated with dissolution of fibrous tissue (*MMP2* and *MMP9*) in myocardium following HLR (myocardium with reduced fibrosis; High*E*+TGFβ→Low*E* substrate). *a*, *n=*2 biologically independent samples per culture condition. *b*, *n=*6 biologically independent samples per culture condition; mean±s.e.m. values are shown. \**P*<0.05 versus Low*E*→Low*E*; ^#^*P*<0.05 versus High*E*+TGFβ→High*E*+TGFβ by one-way ANOVA. *c*, scale bars: 50 μm. *n=*4 biologically independent samples per culture condition; mean±s.e.m values are shown. \**P*<0.05, \*\**P*<0.01, \*\*\**P*<0.005, \*\*\*\**P*<0.001 versus the other group(s) by one-way ANOVA; *d*, *n=*6 biologically independent samples per culture condition; mean±s.e.m. values are shown. *e*, *f*, *n=*4 biologically independent samples per culture condition; mean±s.e.m. values are shown;\**P*<0.05 Vehicle versus Z-DEVD-FMK; ^#^*P*<0.05 versus Low*E*→Low*E*; ^‡^*P*<0.05 versus Low*E*→Low*E*+Z-DEVD-FMK; ^†^*P*<0.05 versus High*E*+TGFβ→High*E*+TGFβ; ^§^*P*<0.05 versus High*E*+TGFβ→High*E*+TGFβ+Z-DEVD-FMK, all by one-way ANOVA.

**Fig. 3.**
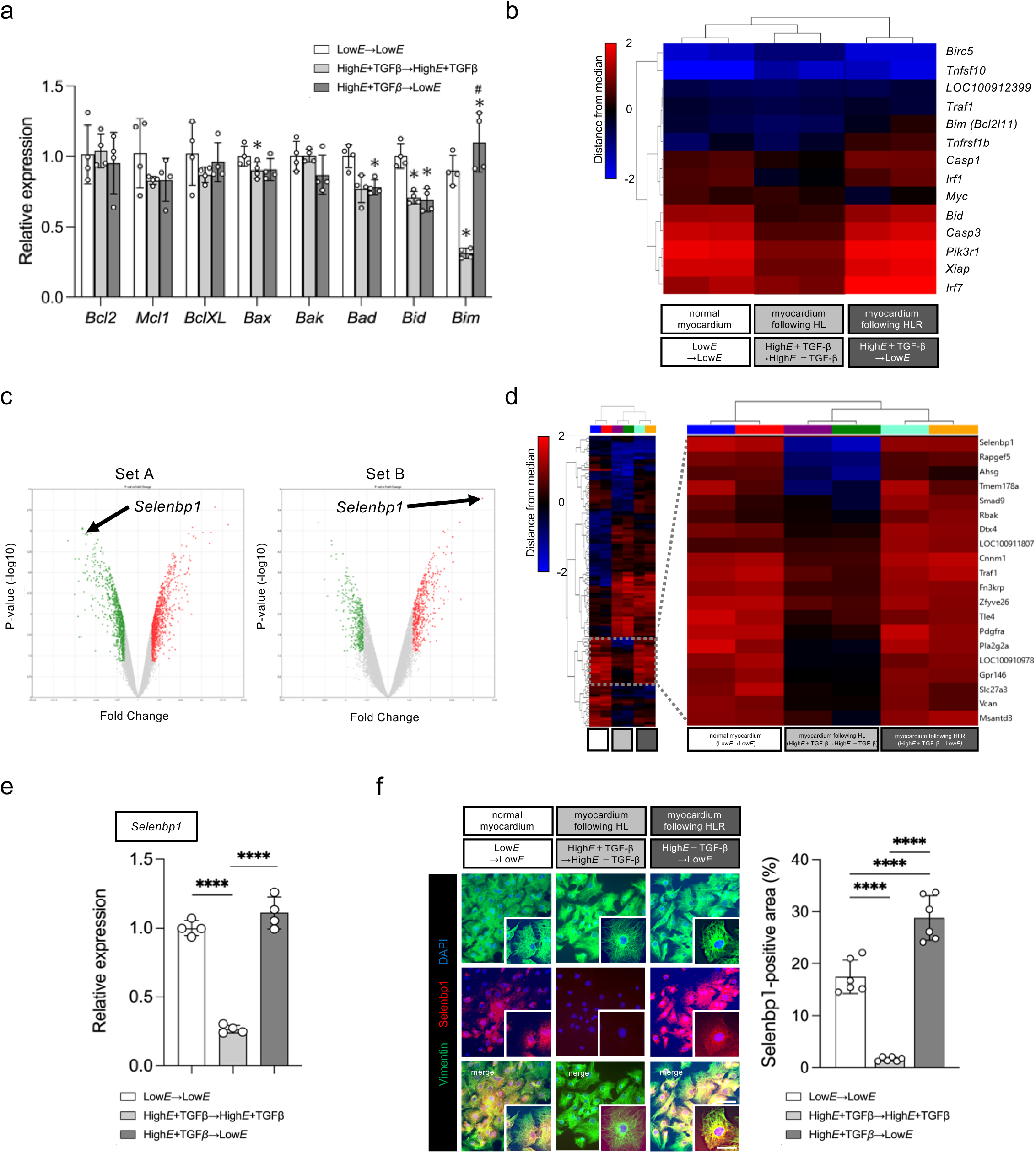
*Selenbp1* is one of the molecules that regulate apoptosis of cultured cardiac fibroblasts via an intrinsic pathway. *a,* qRT-PCR revealed significant downregulation of proapoptotic gene (i.e., *Bim*) in myocardium following HL (fibrotic myocardium; High*E*+TGFβ→ High*E*+TGFβ substrate). *Bim* expression was significantly upregulated in myocardium following HLR (myocardium with reduced fibrosis; High*E*+TGFβ→Low*E* substrate). *b,* Comprehensive gene expression analysis showed significant changes in expression of 14 genes associated with the apoptotic signaling pathway, including *Bim*. *c,* Position of *Selenbp1* on the volcano plots showing gene expression in Set A and Set B. Red indicates genes whose expression was significantly upregulated, and green indicates genes whose expression was significantly downregulated. *d,* A group of genes whose expression was significantly downregulated in myocardium following HL and significantly upregulated in myocardium following HLR compared to expression levels in normal myocardium was extracted. Of these genes with significant changes in expression, *Selenbp1* showed the greatest change in expression. *e*, qRT-PCR revealed significant downregulation of *Selenbp1* in myocardium following HL. Expression was restored in myocardium following HLR. *f,* Representative images of cardiac fibroblasts stained immunocytochemically for Selenbp1. Selenbp1 expression was markedly decreased in myocardium following HL, whereas Selenbp1 expression was markedly increased in myocardium following HLR. *a, n=*4 biologically independent samples per culture condition; mean±s.e.m. values are shown. \**P*<0.05 versus Low*E*→Low*E*; ^#^*P*<0.05 versus High*E*+TGFβ→High*E*+TGFβ, both by one-way ANOVA. *e*, *n=*4 biologically independent samples per culture condition; mean±s.e.m. values are shown. \**P*<0.05, \*\**P*<0.01, \*\*\**P*<0.005, \*\*\*\**P*<0.001 versus the other condition(s), all by one-way ANOVA; *f*, Scale bars: 50 μm (low magnification), 10 μm (high magnification). *n=*6 biologically independent samples per culture condition; mean±s.e.m. values are shown, \**P*<0.05, \*\**P*<0.01, \*\*\**P*<0.005, \*\*\*\**P*<0.001 versus the other condition(s), all by one-way ANOVA.

### Selenbp1 regulates apoptosis of cardiac fibroblasts via an intrinsic pathway

Apoptosis of fibroblasts may involve either an extrinsic or intrinsic pathway (Supplementary Fig. 4a). In our *in vitro* model, death ligands (i.e., tumor necrosis factor alpha [TNFα] and Fas ligand [FasL]) were supplied only by fibroblasts; no other cell type was included. Furthermore, gene expression analysis showed little variation in the expression of *TNFα* and *FasL* and their receptor genes in cultured fibroblasts (Supplementary Fig. 4b, c), suggesting improbability of apoptosis via an autocrine extrinsic pathway. In addition, analysis of enzymatic activity of activated caspase in cultured fibroblasts showed no significant change in caspase-8 or -10 related to the extrinsic pathway (Fig. 2b). Thus, it seemed unlikely that the extrinsic pathway of apoptosis was implicated in our *in vitro* model. qRT-PCR-based analysis of Bcl-2-related genes that regulate the intrinsic pathway of apoptosis showed expression of *Bim* (encoding Bcl-2-like protein 11 [Bcl2l11], a pro-apoptosis protein), to be significantly decreased in myocardium following HL and significantly re-elevated in myocardium following HLR (Fig. 3a). Microarray-based pathway analysis of cultured fibroblasts also showed *Bim* to be one of the apoptosis-related genes that changed significantly in expression in response to changes in the pericellular environment (Fig. 3b). The *Bim* expression patterns matched the variation in caspase-3/7 activity (Fig. 2b, c). To identify molecules that upregulate Bim in this model, bioinformatics analysis was performed based on results of the microarray. We thus focused on *Selenbp1*, which showed the same trend in expression pattern as that of *Bim*, i.e., the greatest change in expression among the genes that were down-regulated in Set A and up-regulated in Set B (Fig. 2c). As shown in the heatmap (Fig. 4d), *Selenbp1* ranked highest among the 20 genes with the same expression pattern as that of *Bim*. Analysis based on GeneMANIA (http://genemania.org) showed that intimate association between *Selenbp1* and a group of apoptosis-related genes has not yet been revealed compared to the close association between *Bim*, *caspase-3*, and apoptosis-related genes (Supplementary Fig. 4d). qRT-PCR showed *Selenbp1* to be significantly downregulated in myocardium following HL and re-elevated in myocardium following HLR (Fig. 3e). The result of immunocyte staining was consistent with this finding (Fig. 4f). Thus it appeared that fibroblast apoptosis in our experimental system was regulated via an intrinsic pathway.

**Fig. 4.**
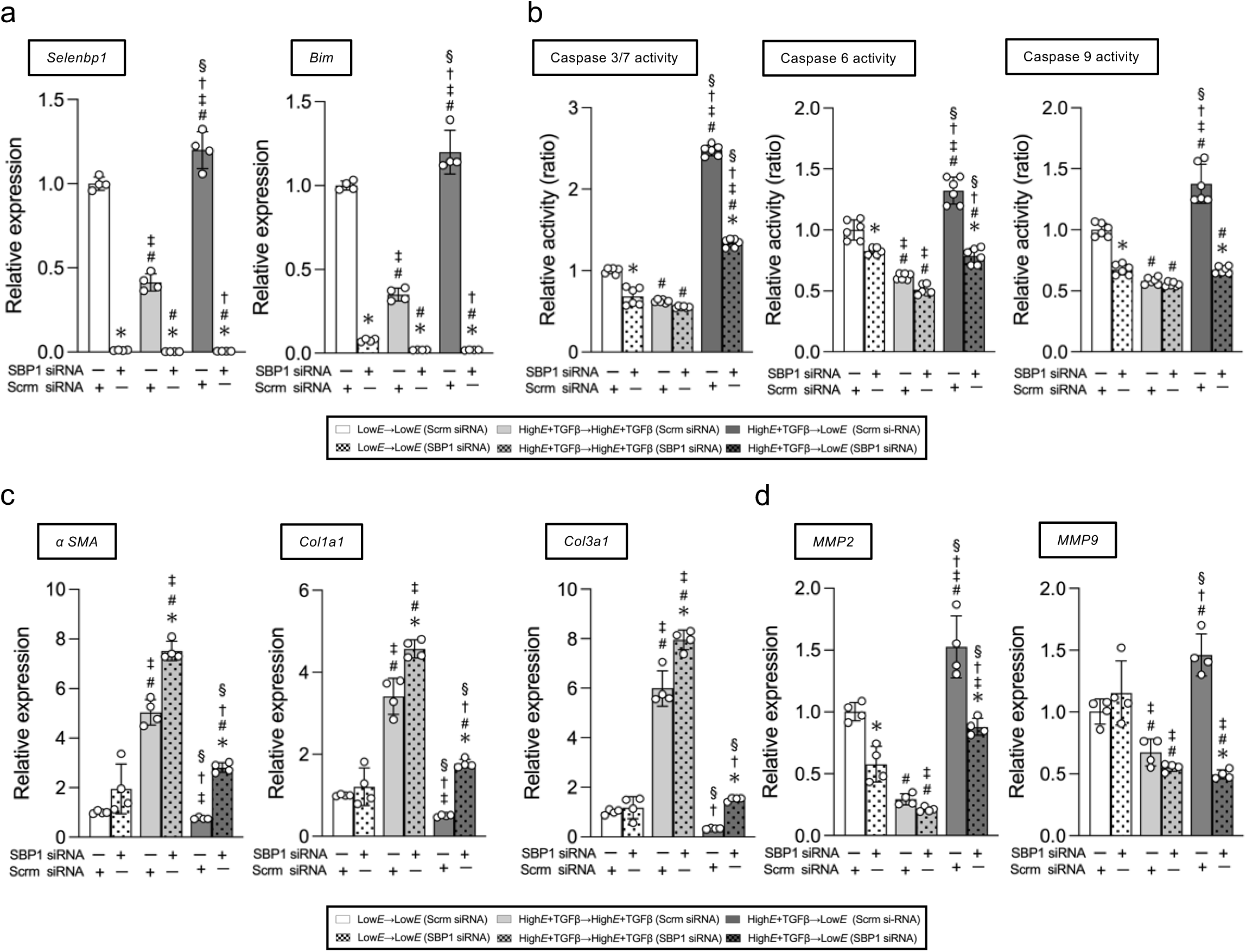
*Selenbp1* knockdown promotes cultured cardiac fibroblast activation and attenuates their differentiation into the fibrinolytic cells. *a,* qRT-PCR revealed both *Selenbp1* and *Bim* expression to be markedly decreased by transfection of Selenbp1 (SBP1) siRNA. *b,* Transfection of SBP1 siRNA markedly decreased caspase3/7, -6, and -9 enzyme activity, especially in the High*E*+TGFβ→Low*E* condition. *c,* qRT-PCR revealed transfection of SBP1 siRNA significantly upregulated fibrosis-associated genes (*αSMA, Col1a1,* and *Col3a1*) in fibroblasts in myocardium following HL (fibrotic myocardium; High*E*+TGFβ→High*E*+TGFβ substrate). *d,* qRT-PCR revealed that transfection of *SBP1* siRNA into fibroblasts significantly downregulated genes-associated with dissolution of fibrous tissue (*MMP2* and *MMP9*) in following HLR (myocardium with reduced fibrosis; High*E*+TGFβ→Low*E* substrate). SBP1, Selenbp1; *a*, *c*, *d*, *n=*4 biologically independent samples per culture condition; mean±s.e.m. values are shown; *b*, *n=*6 biologically independent samples per culture condition; mean±s.e.m. values are shown. \**P*<0.05 scrm siRNA versus SBP1 siRNA, ^#^*P*<0.05 versus Low*E*→Low*E* (scrm siRNA), ^‡^*P*<0.05 versus Low*E*→Low*E* (SBP1 siRNA), ^†^*P*<0.05 versus High*E*+TGFβ→High*E*+TGFβ (scrm siRNA), ^§^*P*<0.05 versus High*E*+TGFβ→High*E*+TGFβ (SBP1 siRNA), all by one-way ANOVA.

### Selenbp1 knockdown promotes activation of cardiac fibroblasts into myofibroblasts and inhibits their differentiation into fibrinolytic forms

To confirm direct contribution of Selenbp1 to the regulation of fibroblast differentiation, we knocked down *Selenbp1* in fibroblasts under each culture condition (Supplementary Fig. 5a) and confirmed gene expression and phenotypic change in the fibroblasts. Transfection of *Selenbp1* siRNA into cultured fibroblasts confirmed a significant decrease in *Selenbp1* and *Bim* expression (Fig. 4a) and suppression of caspase3/7, -6, and -9 activities, which are related to the intrinsic pathway of apoptosis (Fig. 4b). Selenbp1 knockdown significantly increased the expression of fibrosis-related genes (*αSMA, Col1a1, Col3a1*) in myocardium following HL and following HLR (Fig. 4c), and this result was similar to results of quantified immunocyte staining (Supplementary Fig. 5b). However, the expression of fibrinolysis-related genes (*MMP2, MMP9*) and the proportion of immunocyte staining were significantly reduced by Selenbp1 knockdown (Fig. 4d, Supplementary Fig. 5c). Inhibition of *Selenbp1* expression in cultured fibroblasts enhanced fibroblast activation in myocardium following HL and prevented myofibroblast deactivation and differentiation to fibrinolytic cells in myocardium following HLR. These results indicate that Selenbp1 may be one of the molecules that regulate fibroblast differentiation through regulation of the intrinsic pathway of apoptosis in response to the pericellular environment.

### Selenbp1 knockdown attenuates left ventricular contractility and exacerbates diastolic dysfunction due to excessive fibrosis

To elucidate the regulatory mechanism of myocardial collagen turnover in myocardial environments with different hemodynamic loads, we established a normal heart condition (normal rat heart), a pressure-loaded heart condition by subjecting hearts to TAC (TAC-subjected heart), and a reduced pressure-loaded heart condition with TAC removed 4 weeks after TAC (heart released from TAC). Within each group, subgroups of cells transfected with scrambled (scrm) siRNA or *Selenbp1* siRNA were created and compared (Fig. 5a). In scrm siRNA-transfected groups, M-mode echocardiography revealed a significant increase in left ventricular wall thickness and a reduction in left ventricular volume of TAC-subjected heart compared to those of normal heart. These alterations improved with the reduction in afterload. Left ventricular ejection fraction (LVEF) and fractional shortening, indicators of left ventricular contractility, did not change significantly between groups due to compensatory left ventricular hypertrophy (Fig. 5b, c, Supplementary Fig. 6a). *Selenbp1* knockdown in normal heart did not significantly alter left ventricular wall thickness, inner diameter, or contractility compared to those in normal heart transfected with scrm siRNA, whereas *Selenbp1* knockdown in TAC-subjected heart caused a significant increase in left ventricular wall thickness and decrease in contractility. In the heart released from TAC with *Selenbp1* knockdown, there was no significant improvement in these echocardiographic parameters even after the reduction in afterload (Fig. 5b, c, Supplementary Fig. 6a). Evaluation of hemodynamic variables via pressure measuring catheter showed an increase in end diastolic pressure (EDP), the duration of diastole, and the time constant (tau) of left ventricular pressure, reflecting diastolic dysfunction due to myocardial hypertrophy, in TAC-subjected heart compared to normal heart. These hemodynamic variables improved in heart released from TAC (Fig. 5d, e). *Selenbp1* knockdown in normal hearts did not significantly alter hemodynamic parameters, whereas Selenbp1 knockdown in TAC-subjected hearts resulted in reduced contractility and progressive diastolic failure. In heart released from TAC with *Selenbp1* knockdown, there was no significant improvement in these hemodynamic variables even after a reduction in afterload (Fig. 5d, e). We then investigated the causes of these echocardiographic and hemodynamic changes in terms of histology. Extracted heart weight corrected for body weight and body size showed the same trend as left ventricular wall thickness measured by M-mode echocardiography (Supplementary Fig. 6b). A significant increase in left ventricular wall thickness was observed in the TAC-subjected heart (short axis view) compared to that in normal heart, which was further increased in TAC-subjected heart with *Selenbp1* knockdown. Left ventricular wall thickness improved to the same level as that of normal heart in heart released from TAC, but there was no reduction in left ventricular wall thickness in heart released from TAC with *Selenbp1* knockdown (Fig. 5f). In TAC-subjected heart, collagen deposition was observed mainly in the interstitial and periarterial area, and the amount of deposition was significantly increased in the TAC-subjected heart with *Selenbp1* knockdown. Collagen deposition in the interstitial and peri-arterial areas was decreased in heart released from TAC, but no significant decreases were seen in heart released from TAC and subjected to *Selenbp1* knockdown (Fig. 5g). Perivenous collagen deposition did not differ significantly between groups with or without *Selenbp1* knockdown compared to normal heart (Fig. 5g). Expression levels of *Col1a1* and *Col3a1* in cardiac fibroblasts collected from heart reflected the amount of collagen deposition in the interstitial and periarterial areas (Fig. 5h).

**Fig. 5.**
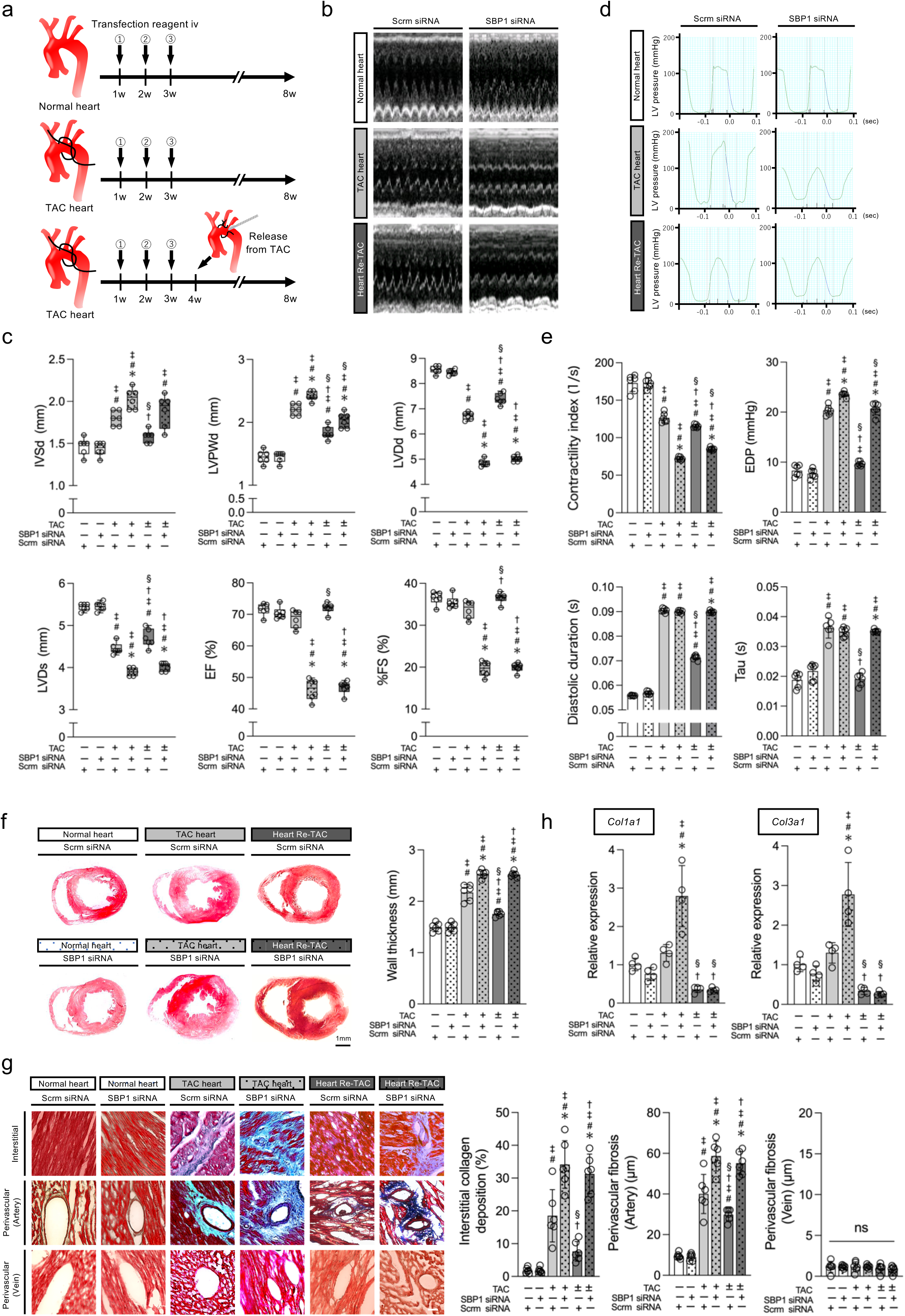
*In vivo Selenbp1* knockdown promotes myocardial fibrosis and exacerbates left ventricular diastolic dysfunction. *a*, Schematic representation of our *in vivo* experiment. *Selenbp1* (*SBP1*) siRNA or scrm siRNA combined with JetPEI^®^ was injected into the tail vein of rats 1 week after sham surgery (normal heart) or after transverse aortic constriction (TAC) (TAC-subjected heart), followed by a total of three intravenous infusions (one given every other week). TAC release was performed 4 weeks after the TAC surgery, reducing afterload (Heart Re-TAC). *b,* Representative M-mode echocardiography tracings of normal heart, TAC-subjected heart and Heart Re-TAC transfected with scrm or *SBP1* siRNA. *c,* Echocardiography revealed increased left ventricular wall thickness and impaired contractility in TAC-subjected heart and improvement in such changes Heart Re-TAC. *SBP1* knockdown further increased left ventricular wall thickness and impaired contractility in TAC-subjected heart, and it inhibited the improvement of these variables in Heart Re-TAC. *d,* Representative left ventricular pressure waveforms in normal heart, TAC-subjected heart, and Heart Re-TAC subjected to scrm or *SBP1* siRNA transfection. *e,* Cardiac catheterization revealed impaired contractility and diastolic dysfunction in TAC-subjected heart but improvement in Heart Re-TAC. *SBP1* knockdown further impaired contractility and increased EDP in TAC-subjected heart but inhibited improvement of these changes in Heart Re-TAC. *f,* Thickness of the left ventricular free wall in short-axis sections of myocardium. TAC increased wall thickness, and release from TAC reduced the increase. *SBP1* knockdown contributed to hypertrophy and prolonged the hypertrophy even after release from TAC. *g,* Masson’s trichrome staining showed increased collagen deposition in the interstitial and peri-arterial area of TAC-subjected heart. *SBP1* knockdown contributed to the increased collagen deposition in TAC-subjected heart and even in Heart Re-TAC. Collagen deposition in peri-venous areas was less than that in the interstitial and peri-arterial areas. *h,* qRT-PCR demonstrated increased expression of *Col1a1* and *Col3a1* in TAC-subjected heart and decreased expression in Heart Re-TAC. *SBP1* knockdown contributed to the increased expression in TAC-subjected heart. TAC ±, surgery to remove TAC 4 weeks after initial TAC surgery; IVSd, interventricular septal thickness at diastole; LVPWd, left ventricular posterior wall thickness at diastole; LVDd, left ventricular diastolic dimension; LVDs, left ventricular systolic dimension; EF, ejection fraction; FS, fractional shortening; Tau, left ventricular diastolic time constant; *c*, *e-g*, *n*=6 animals per group; mean±s.e.m. values are shown. *h*, *n*=4 animals per group; mean±s.e.m. values are shown. \**P*<0.05 scrm siRNA versus SBP1 siRNA; ^#^*P*<0.05 versus normal heart (scrm siRNA), ^‡^*P*<0.05 versus normal heart (*SBP1* siRNA), ^†^*P*<0.05 versus TAC-subjected heart (scrm siRNA), ^§^*P*<0.05 versus TAC-subjected heart (*SBP1* siRNA), all by one-way ANOVA.

### In vivo Selenbp1 knockdown promoted activation of cardiac fibroblasts to myofibroblasts and inhibited differentiation to fibrinolytic cells

qRT-PCR and histological analysis showed that expression of *Selenbp1* and of *Bim* was significantly decreased in fibroblasts collected from TAC-subjected hearts and increased in fibroblasts collected from heart released from TACs compared to normal hearts (Fig. 6a), demonstrating that the expression patterns of *Selenbp1* and *Bim* were similar to that in the *in vitro* pathological model (Fig. 4a). Efficacy of *Selenbp1* siRNA transfection was validated by qRT-PCR. Expression of *Selenbp1* and *Bim* in fibroblasts was decreased in Selenbp1 knockdown groups (Fig. 6a). Caspase-3/7 activity in hearts transfected with scrm siRNA was significantly decreased in TAC-subjected hearts and increased in hearts released from TACs compared to normal hearts (Fig. 6c). Caspase-3/7 activity in hearts transfected with *Selenbp1* siRNA was further decreased in TAC-subjected hearts and declined in the opposite direction in hearts released from TAC (Fig. 6c), trends similar to those in the *in vitro* model (Fig. 4b). The number of vimentin-positive fibroblasts was increased in TAC-subjected hearts compared to normal hearts and decreased in heart released from TAC, and Selenbp1 knockdown increased the number of accumulating fibroblasts (Fig. 6b). The number of αSMA^+^ myofibroblasts was significantly increased in TAC-subjected hearts, and decreased in hearts released from TAC, and Selenbp1 knockdown accelerated accumulation of myofibroblasts in TAC-subjected hearts (Fig. 6d). These trends were supported by qRT-PCR of *αSMA* in fibroblasts (Fig. 6e). The number of MMP2^+^ and/or MMP9^+^ fibrinolytic cells was significantly increased in hearts released from TAC and decreased in hearts released from TAC. Selenbp1 knockdown decreased fibrinolytic cells in heart released from TAC (Fig. 6f). These trends were also supported by qRT-PCR of fibrinolysis-related genes (*MMP2, MMP9*) in fibroblasts (Fig. 6g). Briefly, Selenbp1 knockdown promoted fibroblast activation in TAC-subjected heart (Fig. 6d, e) and inhibited their differentiation into fibrinolytic cells in heart released from TAC (Fig. 6f, g). The above results suggest that Selenbp1 is associated with the regulation of fibroblast differentiation in TAC-subjected heart and heart released from TAC.

**Fig. 6.**
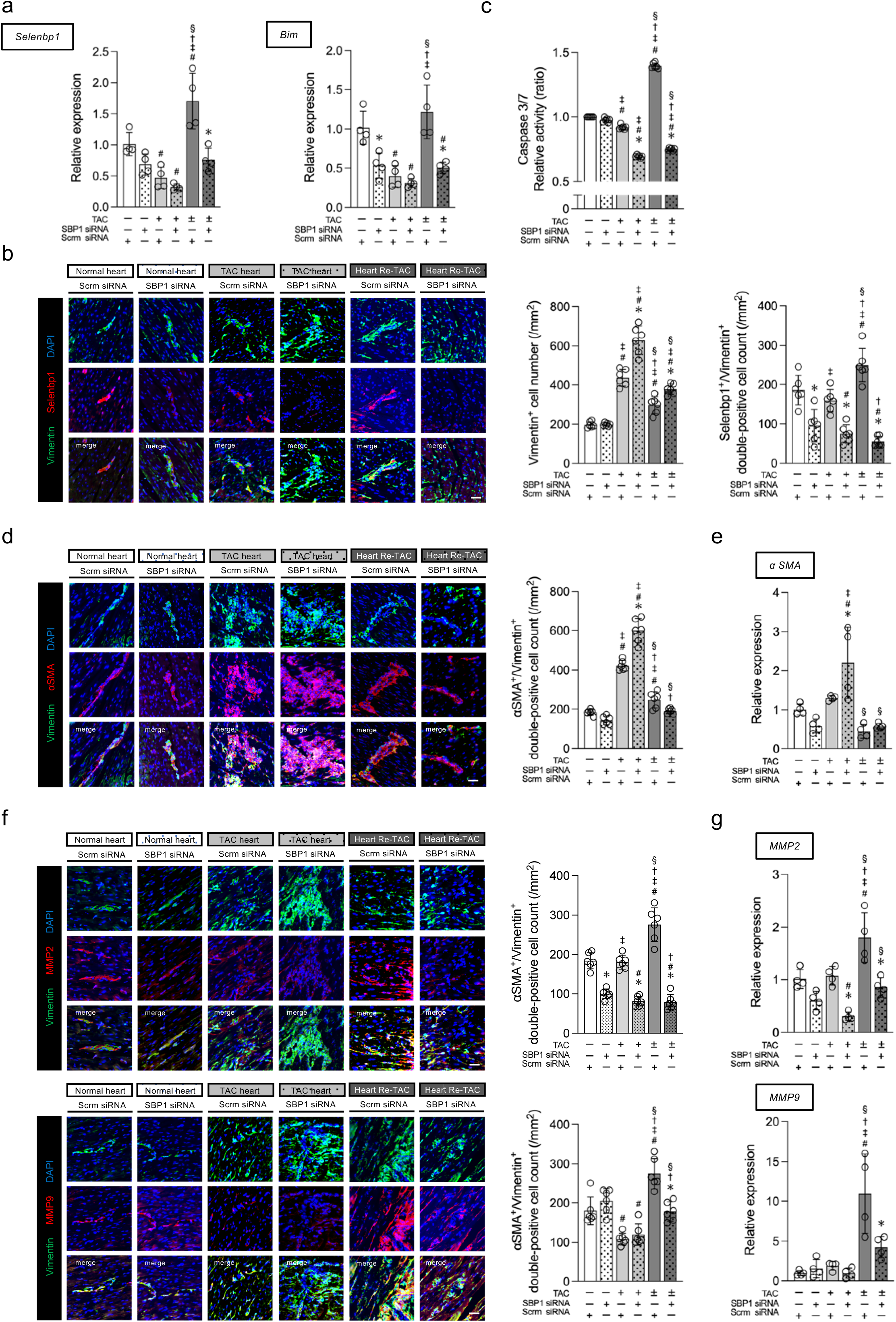
*In vivo Selenbp1* knockdown accelerates cardiac fibroblasts activation and attenuates differentiation of these fibroblasts into fibrinolytic cells. *a,* qRT-PCR revealed decreased expression of *Selenbp1* and *Bim* in transverse aortic constriction (TAC)-subjected heart and increased expression in Heart Re-TAC. *SBP1* knockdown contributed to the decreased expression. *b*, Double immunofluorescence staining showed the number of vimentin-positive cells to be increased in TAC-subjected heart and decreased in Heart Re-TAC. *SBP1* knockdown contributed to the increased number of vimentin-positive cells in TAC-subjected heart and in Heart Re-TAC. The number of vimentin- and Selenbp1 double-positive cells was increased in Heart Re-TAC. *SBP1* knockdown contributed to a decrease in the number of vimentin- and Selenbp1 double-positive cells in all groups. *c,* Enzyme activity of caspase-3/7 significantly increased in Heart Re-TAC. *SBP1* knockdown contributed to the decrease in these enzyme activities. *d,* Double immunofluorescence staining revealed an increase in the number of vimentin and αSMA double-positive cells in TAC-subjected heart and a decrease in Heart Re-TAC. *SBP1* knockdown contributed to further increase the number of vimentin- and αSMA double-positive cells in TAC-subjected heart. *e,* qRT-PCR revealed increased expression of *αSMA* in TAC-subjected heart and decreased expression in Heart Re-TAC. *SBP1* knockdown contributed to the increased expression in TAC-subjected heart. *f,* Double immunofluorescence staining revealed an increase in vimentin- and MMP2 or MMP9 double-positive cells in Heart Re-TAC. *SBP1* knockdown contributed to a decrease in the number of vimentin- and MMP2 or MMP9 double-positive cells in Heart Re-TAC. *g,* qRT-PCR revealed increased expression of *MMP2* and *MMP9* in Heart Re-TAC. *SBP1* knockdown contributed to a decrease in *MMP2* and *MMP9* expression in Heart Re-TAC. TAC±, surgery to remove TAC 4 weeks after initial TAC surgery. *b, d, f,* Scale bars: 100 μm. *a, e, g*, *n*=4 animals per group and *b, c, d*, *f*, *n*=6 animals per group; mean±s.e.m. values are shown; \**P*<0.05 scrm siRNA versus SBP1 siRNA, ^#^*P*<0.05 versus normal heart (scrm siRNA); ^‡^*P*<0.05 versus normal heart (*SBP1* siRNA); ^†^*P*<0.05 versus TAC-subjected heart (scrm siRNA); ^§^*P*<0.05 versus TAC-subjected heart (*SBP1* siRNA), all by one-way ANOVA.

## Discussion

The microenvironment, defined as the extracellular space surrounding and supporting the cell, is composed of the ECM, soluble chemical factors and adjacent cells. Fibroblasts play an important role as the primary cells producing the ECM that maintains the layered structure of the myocardium, and are not only a passive scaffold, but are essential for the regulation of normal cellular function and the maintenance of organ homeostasis^19, 20^. It has been shown that valvular stromal cells, the major fibroblast population in heart valves, are quiescent in normal valves with a tissue elastic modulus of about 0.8-8 kPa, but differentiate into myofibroblasts and osteoblast-like cells in pathological valves with progressive valve stenosis and a tissue elastic modulus as stiff as 27±10 kPa^21–23^, and this phenotype has been reported to revert when the substrate elastic modulus is again reduced to normal^24, 25^. However, the pathways by which myofibroblasts are deactivated in the heart and the potential molecular mechanisms by which myofibroblasts transition to fibrinolytic cells are not understood. One reason for this is the lack of appropriate *in vitro* disease models to fully understand the cellular and molecular basis of myocardial fibrosis. A unique feature of this study was the *in vitro* reproduction of a pericellular environment that mimics normal myocardium, fibrotic myocardium following hemodynamic loading, and even myocardium with reduced fibrosis following hemodynamic loading reduction. Although plastic substrates for cell culture have become the traditional substrate for mammalian cell culture because they support basic cell survival and function, the elastic modulus of most cell culture plastics exceeds 1 GPa, many orders of magnitude stiffer than the natural growth environment of cardiac fibroblasts. Mechanical stress from culture at high elasticity may significantly alter the genomic transcriptional program and affect the spontaneous activation of cardiac fibroblasts. Previously, there has been a focus on modulating the elastic modulus of the substrate to avoid the problem of spontaneous activation under standard cell culture conditions^26^. In the present study, we utilized silicone gel as a culture substrate to better mimic the ECM environment of myocardium following HL. These are culture media with a thin layer of chemically functionalized, stable, modulus-verified biocompatible silicone that form covalent bonds with the amines on the proteins at the bottom of the culture well. Silicon gels, unlike hydrogels, do not undergo hydrolysis, do not dry out or swell, and can be adjusted to various elastic moduli. Although silicone gels do not perfectly mimic the native ECM environment, they appeared to better recapitulate the normal and fibrotic myocardial niches that influence the phenotype of cardiac fibroblasts. The elastic modulus of the normal adult rat heart is about 18±2 kPa or 11.9 to 46.2 kPa, and the elastic modulus of rat myocardium with advanced fibrosis is reported to be 55±15 kPa^27, 28^. Considering these reports, the present study employed a silicone gel substrate with an elastic modulus of 16 kPa for normal myocardium and 64 kPa for myocardium following HL. In addition, TGFβ plays the most significant role in pathological fibrosis, and the persistence of fibroblast activation is considered to be a consequence of the persistence of TGFβ signaling^29, 30^. TGFβ was added to the medium of fibrotic myocardium (i.e., myocardium following HL) in this study. TGFβ expression is known to be decreased in myocardium following HLR^31^. Therefore, addition of TGFβ was omitted in myocardium following HLR. *In vitro* expression levels of αSMA, Col1a1, and Col3a1 as well as results of quantified immunostaining in our normal myocardium and myocardium following HL showed the same trends as *in vivo* gene expression levels and immunostaining results in our normal and myocardium following HL with TAC, supporting the notion that 16 kPa and 64 kPa silicone gels with TGFβ can be used as substrates for *in vitro* models that recapitulate normal and fibrotic myocardial environments.

In this study, we demonstrated that cardiac fibroblasts sense and respond to changes in the elasticity of the culture substrate. Substrate stiffening induced a decrease in caspase-3 activity and fibroblast activation, whereas the change from substrate stiffening to softening led to an increase in caspase-3 activity led to fibroblast deactivation and phenotypic change to fibrinolytic type. These results indicate that the apoptotic signaling pathway is associated with differentiation of fibroblasts in response to the pericellular environment. Anti-apoptotic protein expression induced by ECM stiffness is regulated by YAP and TAZ, and translocation to the nucleus is induced by integrin and FAK-mediated mechanotransduction pathways^32, 33^. These findings and the results of comprehensive gene expression analysis of cultured fibroblasts in this research model suggested that the activation and deactivation of fibroblasts in response to stiffness in culture substrates involves suppression and promotion, respectively, of the apoptotic signaling pathway via the integrin-FAK-ROCK-YAP/TAZ cascade. The PI3K/AKT pathway is also associated with myofibroblast activation on stiff substrates, and endogenous AKT activation correlates with cell phenotype^19^. However, there was no significant change in the PI3K/AKT signaling pathway in this study model. The soluble chemical factor TGFβ, a component of the microenvironment, also induces integrin expression in cardiac fibroblasts via the TGFβ-SMAD pathway^34^. The TGFβ signaling pathway showed significant variation in our comprehensive expression analysis. TGFβ recombinant protein was added *in vitro* to myocardium following HL. Therefore, integrin expression may have been induced via the TGFβ-SMAD pathway, resulting in suppression of the apoptotic signaling pathway. Other fibroblast activation pathways include induction of the cardiac fibrosis pathway by the RhoA/ROCK pathway^35, 36^, fibroblast activation caused by oxidative stress-induced activation of the ROS system^37^, activation of the p38α MAPK pathway^38–40^, and induction of cardiac fibroblast to myofibroblast conversion by activation of the Wnt pathway^41^. However, these known fibroblast activation pathways were not significantly altered in our three-part *in vitro* model.

The molecular basis of regulation of apoptosis of fibroblasts is considered to lie in increased mitochondrial apoptotic priming^32, 42^. The intrinsic pathway of apoptosis in myocardium following HL was inhibited via transcriptional repression of the apoptosis-promoting molecule Bim (Bcl2l11), which could suppress mitochondrial priming in myofibroblasts. However, in myocardium following HLR, the intrinsic apoptosis pathway was promoted via transcriptional enhancement of Bim (Bcl2l11), which could be interpreted as an increase in mitochondrial priming in myofibroblasts leading to apoptotic progress. These findings and the results of our study support the notion that biomechanical signals induced by ECM stiffness regulate apoptosis of fibroblasts via endogenous pathways.

ECM stiffness promotes differentiation of fibroblasts into myofibroblasts, whereas release from stiffness results in loss of contractile function and reduced activation of myofibroblasts^43, 44^. In our previous study evaluating fibroblast differentiation and apoptosis and substitutive fibrosis in tissue repair after myocardial infarction, we observed fibroblast activation and suppression of apoptosis in tissues with increased substitutive fibrosis and sclerosis^45^, suggesting a link between phenotypic regulation and apoptosis of cardiac fibroblasts. An important finding of our study is that activation of the apoptotic signaling pathway is associated with myofibroblast deactivation and their differentiation into fibrinolytic cells during the transition from fibrotic myocardium to myocardium with reduced fibrosis.

This study focused on biochemical mediators that can regulate fibroblast apoptosis and alter the phenotype of fibroblasts. The human Selenbp1 protein is ubiquitously expressed in a variety of tissues and is highly expressed in heart, lung, liver, and kidney^46^. Since the first report of Selenbp1 as a tumor-associated protein in prostate cancer^47^, decreased expression or loss of Selenbp1 has been observed in various solid tumors^48^. On the other hand, increased expression of Selenbp1 has been shown to significantly suppress cancer cell malignancy^49^. These findings shed light on the mechanism by which Selenbp1 knockdown led, in our study, to suppression of fibroblast apoptosis, resulting in increased ECM deposition by promoting fibroblast activation and decreased ECM degradation by suppressing conversion to fibrinolytic cells, worsening of myocardial fibrosis.

Cells involved in tissue repair after myocardial injury include fibroblasts, endothelial cells, pericytes, and immune cells. Because we attempted to elucidate, *in vivo*, the relative contribution of fibroblasts in myocardium following HL and HLR, our analysis did not incorporate soluble chemical factors, cardiomyocytes, macrophages, or endothelial cells, which are in their native niche, and did not account for the complex intercellular network in fibrosis. Further clarification is needed, so a model that more closely resembles the *in vivo* environment must be developed. Furthermore, in addition to fibroblasts derived from cells of embryonic epicardium, fibroblasts derived from vascular endothelial cells^50^, perivascular cells^51^, hematopoietic bone marrow-derived progenitor cells^52, 53^, and circulating fibroblasts^54^ have also been reported to contribute to pathological cardiac remodeling. Our three-part *in vitro* model, in which we used resident fibroblasts derived from cells of embryonic epicardium, also differs from the *in vivo* environment in that it does not consider alternative sources of these activated fibroblast populations. In addition, *in vivo* transfection using nanoparticles resulted in knockdown of Selenbp1 in an unspecified number of cells; studies incorporating cardiac fibroblast-specific Selenbp1 knockout rats are needed to clarify the role of Selenbp1.

The findings of this study on cardiac fibroblasts provide insight into some of the cellular and molecular mechanisms that regulate ECM degradation, a key event in the process of fibrosis reversal. Control of fibroblast apoptosis could become a new therapeutic strategy to reverse established fibrosis not only by inducing deactivation of myofibroblasts but also by inducing their differentiation to fibrinolytic cells. Although further efforts are needed to understand the resolution of fibrosis, this study, which elucidated one aspect of the regulatory mechanism of differentiation of cardiac fibroblasts, may lead to new treatment strategies for patients with heart failure and associated myocardial fibrosis, the incidence of which is on the rise. Thus, our findings have important implications for improving the prognosis of heart failure.

## Methods

### Isolation of cardiac fibroblasts

Cardiac fibroblasts were isolated from the hearts of rats, as described previously^55, 56^ but with some modifications. Eight-week-old male Sprague-Dawley rats were obtained from The Jackson Laboratory and housed under standard conditions (controlled light/dark cycle, temperature, and humidity) in cages at the animal facility of Saitama Medical Center in Saitama, Japan. The hearts were cut into 1-mm^3^ fragments, and when cultured were evenly spaced with none touching another. Fragments of normal myocardium were placed in 0.1% gelatin-coated CytoSoft 16 kPa tissue culture dishes (Advanced Biomatrix). Each fragment was covered with a drop of Dulbecco’s Modified Eagle Medium (DMEM) (Sigma-Aldrich) containing 2% FBS, 50 U/ml penicillin, and 50 µg/ml streptomycin and incubated for 24 hours at 37°C in a CO2 incubator. Sufficient medium was then added to cover the entire bottom of the culture dish, and the culture was incubated for an additional 24 hours. After this period, sufficient medium was added to completely cover the tissue fragments, and culturing was continued for another 5 days. When fibroblast outgrowth was observed, the cells were collected by trypsinization. These fibroblasts were maintained in DMEM containing 2% FBS, 50 U/ml penicillin, and 50 µg/ml streptomycin in 0.1% gelatin-coated CytoSoft 16 kPa or 64 kPa dishes.

### In vitro representation of cardiac fibrosis

Cardiac fibroblasts (2 × 10^4^) were plated on a 0.1% gelatin-coated CytoSoft dish (Advanced Biomatrix) and cultured for 48 hours in DMEM containing 2% FBS, 50 U/ml penicillin, and 50 µg/ml streptomycin. he cultured fibroblasts were then transferred to a 0.1% gelatin-coated CytoSoft 16 kPa or 64 kPa dish with or without recombinant TGFβ protein (10lng/ml). Forty-eight hours later, the cultured fibroblasts were again transferred to a 0.1% gelatin-coated CytoSoft 16 kPa or 64 kPa dish with or without recombinant TGFβ protein (10lng/ml). In substrate mimicking the elastic modulus of normal myocardium (Low*E*→Low*E* condition), cells were passaged every 48 hours in a low-elasticity substrate (16 kPa). In substrate mimicking the elastic modulus of myocardium following HL (fibrotic myocardium; High*E*+ TGFβ→High*E*+TGFβ substrate), cells were passaged every 48 hours in a high-elasticity substrate (64 kP) with TGFβ. In substrate mimicking the elastic modulus of myocardium following HLR (myocardium with reduced fibrosis; High*E*+TGFβ→Low*E* substrate), cells were incubated in a high-elasticity substrate (64 kPa) with TGFβ for 48 hours, and this was followed by passages in a low-elasticity substrate (16 kPa). The cultured fibroblasts were fixed with 4% paraformaldehyde for immunofluorescence analysis or lysed for mRNA expression analysis.

In vitro *transfection*. Cardiac fibroblasts (2 × 10^4^) were plated on a 0.1% gelatin-coated CytoSoft 6-well dish (Advanced Biomatrix) and cultured for 24 hours in DMEM containing 2% FBS, 50 U/ml penicillin, and 50 µg/ml streptomycin, and transfected with *Selenbp1* siRNA (Invitrogen) with use of RNAiMAX transfection reagent (Invitrogen) in Opti-MEM (Gibco) according to the manufacturer’s protocol. Scrm siRNA sequences were used for negative control. siRNA was used at a final concentration of 40lnM. Scrm siRNA was used for positive control. The transfection mix was added to medium containing all supplements and cultured for 24 hours in DMEM containing 2% FBS, 50 U/ml penicillin, and 50 µg/ml streptomycin. After transfection, fibroblasts were transferred to a 0.1% gelatin-coated CytoSoft 16 kPa or 64 kPa six-well dishes with or without recombinant TGFβ protein (10lng/ml). Forty-eight hours later, the fibroblasts were further transferred to 0.1% gelatin-coated CytoSoft 16 kPa or 64 kPa six-well dishes with or without recombinant TGFβ protein (10lng/ml). The fibroblasts were fixed with 4% paraformaldehyde for immunofluorescence analysis or lysed for mRNA expression analysis.

### RNA extraction and real-time PCR

Total RNA was extracted from cells or heart tissue with a Gene Jet PCR Purification Kit (Thermo Scientific) and quantified with a Nano-Drop 8000 spectrophotometer (Thermo Scientific). For extraction of total RNA from rat hearts, fresh tissue was excised from left ventricular wall. cDNA was synthesized from 150 ng of total RNA extracted from heart tissues with a High Capacity cDNA Reverse Transcription Kit (Applied Biosystems). Real-time PCR was performed with QuantStudio (Applied Biosystems) and SYBR Premix Ex Taq II (Takara Bio, Shiga, Japan) under the following conditions: 95°C for 30 seconds followed by 40 cycles at 95°C for 5 seconds and 60°C for 30 seconds.

Gene expression levels were normalized to *Gapdh* (forward, 5’-CCCCTGGCCAAGGTCATCCA-3’; reverse,

5’-CGGAAGGCCATGCCAGTGAG-3’) expression according to the 2−Δet2−Δct method. Gene expression data were acquired from independent biological replicates. The primers are *αSMA* (forward,

5’-ACAACTGGTATTGTGCTGGACT-3’; reverse,

5’-TGGCATGAGGCAGAGCATAG-3’), *Bad* (forward,

5’-AGGGATGGAGGAGGAGCTTA-3’; reverse,

5’-GGCGAGGAAGTCCCTTGAA-3’), *Bak* (forward,

5’-GCCAGATGGATTGCACAGAG-3’; reverse,

5’-CCACAAATTGGCCCAACAGA-3’), *Bax* (forward,

5’-GGCGATGAACTGGACAACAA-3’; reverse,

5’-ACACGGAAGAAGACCTCTCG-3’), *Bcl2* (forward,

5’-GATAACGGAGGCTGGGATGC-3’; reverse,

5’-CAGGTATGCACCCAGAGTGA-3’), *BclXL* (forward,

5’-AGCCAGAACCCTATCTTGGC-3’; reverse,

5’-GCTCAACCAGTCCATTGTCC-3’), *Bid* (forward,

5’-CTCGGGTCCAAGTGTCGGTC-3’; reverse,

5’-CCTGAGCCATTGCTGACCTC-3’), *Bim* (forward,

5’-TCTGTTCTGGTACGCCACTT-3’; reverse,

5’-GGTCCGAAGAGTTGAGGGAA-3’), *Col1a1* (forward,

5’-TGAGCCAGCAGATTGAGAAC-3’; reverse,

5’-CCAGTACTCTCCGCTCTTCC-3’), *Col3a1* (forward,

5’-AGTCTGGAGTCGGAGGAATG-3’; reverse,

5’-AGGATGTCCAGAGGAACCAG-3’), *FasL* (forward,

5’-TCAGAGGAAACAGTGGGTGC-3’; reverse,

5’-GTTTTGACAGTCCGTTAGGCA-3’), *FasR* (forward,

5’-TAAGCTCCTTTGGCTGCTGA-3’; reverse,

5’-GCACACTTTCAGGACTTGGG-3’), *Mcl1* (forward,

5’-GCTGCATCGAACCATTAGCA-3’; reverse,

5’-GCACATTTCTGATGCCACCT-3’), *Mmp2* (forward,

5’-AGAAGGCTGTGTTCTTCGCA-3’; reverse,

5’-AAAGGCAGCGTCTACTTGCT-3’), *Mmp9* (forward,

5’-TGCCCTGGAACTCACACAAC-3’; reverse,

5’-GTTCACCCGGTTGTGGAAAC-3’), *Selenbp1* (forward,

5’-ACTTGCCCTGCATTTACCGA-3’; reverse,

5’-CAGCCTATGGATGACCTGGC-3’), *TNFR1* (forward,

5’-AGTCTACTGTGCCGATATCCC-3’; reverse,

5’-TCACCCTCCACCTCTTTGAC-3’), *TNFR2* (forward,

5’-TACCCAGGTCTGGAACCATC-3’; reverse,

5’-GTCTCCACCTGGTCATCACT-3’), *TNFα* (forward,

5’-TCGTAGCAAACCACCAAGTG-3’; reverse,

5’-TTGTCTTTGAGATCCATGCC-3’).

### Microarray analysis

Total RNA of was isolated from cardiac fibroblasts as described above and subjected to GeneChip analysis (GeneChip Arrays platform with Clariom™ S Assay, Rat; Affymetrix). Two independent biological replicates were prepared for cardiac fibroblasts. Median per-chip normalization was performed for each array. A fold-change cutoff of 1.5-was used to identify differentially expressed genes.

*Immunocytochemistry*. Cells were fixed in 4% paraformaldehyde/phosphate buffered saline (PBS) for 5 minutes at room temperature. Cells not set aside for surface antigen staining were incubated in PBS containing 0.1% Triton X-100 (X100, Merck) for 5 minutes at room temperature. Non-specific antibody binding sites were pre-blocked with PBS containing 5% goat serum for 30 minutes at room temperature. Primary antibodies were then applied to the cells for 1 hour at room temperature: anti-vimentin Ab at 1:100 dilution (ab24525, Abcam), anti-cleaved caspase 3 Ab at 1:100 dilution (9661, Cell Signaling), anti-αSMA Ab at 1:00 dilution (ab5694, Abcam), anti-collagen1 Ab at 1:100 dilution (ab44710, Abcam), anti-collagen3 Ab at 1:100 dilution (ab7778, Abcam), anti-MMP2 Ab at 1:200 dilution (436000, Invitrogen), anti-MMP9 Ab at 1:200 dilution (MA5-32705, Invitrogen), anti-Selenbp1 Ab at 1:200 dilution (PA5-102394, Invitrogen). Cells were rinsed and then incubated with fluorophore-conjugated secondary antibody at 1:300 dilution (Alexa Fluor 488- or Alexa Fluor 594-conjugated polyclonal antibody; Invitrogen) and 4l,6-diamidino-2-phenylindole (DAPI) in blocking buffer for 1 hour at room temperature. Data were acquired from independent biological replicates.

*Enrichment analysis*. All statistically enriched terms (e.g., GO/KEGG terms, canonical pathways, and hallmark gene sets) were identified, and accumulative hypergeometric *P* values and enrichment factors were calculated and used for filtering. Remaining significant terms were then hierarchically clustered into a tree based on Kappa-statistical similarities among their gene memberships. A kappa score of 0.3 was applied as the threshold to cast the tree into term clusters. The term with the best *P* value within each cluster was selected as the representative enriched term and displayed in a dendrogram. Heatmap cells were color graded according to their *P* values, with white indicating lack of enrichment for a given term in the corresponding gene list.

*Caspase activity assay*. Caspase activity was measured by Caspase Family Fluorometric Substrate Kit II Plus (ab102487, Abcam) according to the manufacturer’s protocol. Briefly, 1 x 10^6^ cultured fibroblasts were suspended in cell lysis buffer and incubated on ice. Reaction buffer and dithiothreitol were added to each sample. AFC-conjugated substrate was added to each sample. After 1–2 hour-incubation at 37°C, samples were read in a fluorometer equipped with a 400-nm excitation filter and 505-nm emission filter.

*TAC and TAC release*. For further experimental validation of bioinformatic findings, cardiac hypertrophy was induced by TAC, as previously described^57^ but with some modifications. After 1 week of adaptation, rats were randomly divided into four groups and underwent sham or TAC surgery with or without siRNA transfection. For TAC, rats (*n*l=l6) were anesthetized with 3-4% isoflurane gas (in 100% oxygen) in an induction chamber and then intubated. Throughout the surgery, anesthesia was maintained with 1.5% isoflurane in 100% oxygen, and body temperature, monitored by rectal thermometer probe, was maintained at 37±l0.5℃ by means of a heat pad. Before anesthesia, 0.1lmg/kg body weight of buprenorphin was injected subcutaneously. The surgical level of anesthesia was confirmed by absence of the toe pinch reflex. The skin of the upper thorax was shaved and disinfected with povidone-iodine and alcohol, and 1lmg/kg bupivacaine was then injected into the upper sternum. Thoracotomy was performed by cutting the upper sternum, and the sternum was retracted with use of a fine retractor to expose the aorta. A piece of 3-0 silk suture was placed around the transverse aorta between the brachiocephalic artery and left carotid artery and tied loosely into a single knot. A presterilized, flat end 21G needle was inserted into the suture knot, which was then tightened fully, encircling the transverse aorta and the blunt-end needle. The suture was secured with a double knot before the needle was removed. The incision was closed in layers with 4-0 suture material. The sham operation included only thoracotomy; there was no constriction (*n*l=l6). The TAC release surgery was performed 4 weeks after the initial TAC surgery under the same anesthesia protocol. Sternotomy was performed by re-incision of the upper sternum. Adhesions were dissected, the knot that had been tied around the aorta was cut, and the compression was released . The incision was closed in layers with 4-0 suture material. Normal hearts and TAC-subjected hearts were collected 8 weeks after the initial surgery., and TAC-subjected hearts were collected 4 weeks the release surgery. Hearts released from TAC were perfused with saline, frozen in liquid nitrogen, and stored at −80l°C. For histologic analysis, hearts were embedded in optimal cutting temperature compound (VWR International).

### In vivo transfection

To investigate the role of Selenbp1 in cardiac fibrosis, we performed *in vivo* gene silencing using Selenbp1 siRNA (AM16830, Invitrogen) along with a transfection reagent, as previously described^58^ but with some modification. Forty micrograms of siRNA combined with the non-viral *in vivo* transfection reagent JetPEI^®^ (Polyplus-transfection) was administered three times every week to each rat through tail-vein injection. An N/P ratio of 8 in a total volume of 200 μL of 5% dextrose was maintained for each injection, following the manufacturer’s protocol. Scrambled siRNA was administered as control.

### Echocardiography

Cardiac function was assessed by means of M-mode transthoracic echocardiography performed with a Vscan Extend R2 system and 8-MHz transducer; GE Healthcare) on day 28 after surgery. Each animal was anesthetized with isoflurane gas, and two-dimensional imaging of the heart was performed and recorded through the anterior and posterior left ventricle (LV) walls. Anterior and posterior wall thicknesses (end-diastolic and end-systolic) and left ventricular internal dimensions were measured during at least three consecutive cardiac cycles, as previously described^59^. To determine left ventricular volume, the maximum minor axis of the left ventricular end-diastolic dimension (LVDd) and left ventricular end-systolic dimension (LVDs) were measured on the parasternal long- or short-axis view, on the assumption that the LV was prolate spheroid in shape. Left ventricular volume was usually calculated by the Teichholz formula: V = 7.0 × D^3^ / (2.4 + D)^60^. Left ventricular stroke volume (SV) was calculated by subtracting left ventricular end-systolic volume (LVESV) from left ventricular end-diastolic volume (LVEDV). LVEF was calculated according to the following formula: LVEF = SV × 100 / LVEDV (%)^60^. Left ventricular fractional shortening (%FS), an index of systolic function, was calculated as follows: %FS = (LVDd − LVDs) × 100 / LVDd (%)^60^. Measurements were obtained at least three times in each 6 animal, and the mean value [each of 6] was used for analysis.

### Measurement of hemodynamic variables

Hemodynamic variables were measured by means of cardiac catheterization, as previously described^61, 62^. Briefly, under general anesthesia achieved with isoflurane and mechanical ventilation, a catheter (PR 671, Millar Instruments) was inserted into the LV cavity through the LV free wall. Intra-left ventricular pressure signals were measured (SPR-671, Millar Instruments) and digitally recorded with a data acquisition system (PowerLab 2/26, ADInstruments). All data were analyzed by LabChart Software (ADInstruments). Measurements were obtained at least five times in each animal, and the mean value [each of 6] was used for analysis. Data were collected from at least five different measurements in a blinded manner.

### Trichrome staining

Frozen tissue sections (8 µm thick) were prepared as described above and stained with Modified Masson’s Trichrome (Trichrome Stain Kit, ScyTek Laboratories) according to the manufacturer’s instructions. Data were acquired from independent biological replicates.

### Immunohistochemistry

Rats were killed by cervical dislocation, and immediately thereafter, the aorta was clamped, and ice-cold PBS was injected into the left ventricular cavity. The heart was then perfused with ice-cold 4% paraformaldehyde in PBS. The heart was removed, cut at the midpoint along the short axis of the left ventricle, embedded in optimal cutting temperature compound (VWR International), and frozen in isopentane chilled in liquid nitrogen. Frozen tissue sections (8 µm thick) were prepared, and non-specific antibody-binding sites were pre-blocked with PBS containing 5% goat serum. Primary antibodies were then applied overnight at 4°C: anti-vimentin Ab at 1:100 dilution (ab24525, Abcam) and anti-αSMA Ab at 1:00 dilution (ab5694, Abcam); anti-MMP2 Ab at 1:200 dilution (436000, Invitrogen), anti-MMP9 Ab at 1:200 dilution (MA5-32705, Invitrogen) and anti-Selenbp1 Ab at 1:100 dilution (PA5-102394, Invitrogen). Cells were rinsed and then incubated with fluorophore-conjugated secondary antibody at 1:300 dilution (Alexa Fluor 488- or Alexa Fluor 594-conjugated polyclonal antibody, Invitrogen) and DAPI in blocking buffer for 1 hour at room temperature. Data were acquired from independent biological replicates.

### Imaging and analysis

Digital images of the trichrome-stained tissue sections were acquired with an all-in-one type fluorescence microscope (BZ-8000, KEYENCE) and imported as TIFF files into ImageJ (National Institutes of Health). The fibrin-positive area of each of five regions was calculated as a percentage of the entire region. The Color Threshold function of ImageJ was applied to measure the fibrin clot area. To measure the fibrin clot area, the image was first divided into three parts by means of the Split Channels function, and the image with the clearest view of the target area was then selected. A threshold was then set by means of the Threshold tool to highlight only the object of interest (by setting the threshold at which only the fibrin clot area was detected) in the image. The percentage of the object in the image was then calculated. The same procedure was used for images of normal heart, TAC-subjected heart, and heart released from TAC with or without Selenbp1 knockdown, and the threshold value set for normal heart was used for heart tissue under the other conditions.

*Statistical analysis*. Statistical analysis was performed with GraphPad Prism version 8 (GraphPad Software). Values were assumed to be normally distributed and are shown as mean±s.e.m. All statistical tests were parametric. For comparison between multiple sample sets, repeated-measures analysis of variance (ANOVA) or one- or two-way ANOVA was performed, followed by Bonferroni’s post-hoc test. For analysis of values between two sample sets, two-tailed, unpaired Student’s *t-*test was used.

## Data availability

The authors declare that all supporting data are available within the article and its online supplementary files. For purposes of reproducibility, additional technical information and data that support the findings of this study are available from the corresponding authors on reasonable request. Microarray data can be obtained from Gene Expression Omnibus GSE214665 (https://www.ncbi.nlm.nih.gov/geo/query/acc.cgi?acc=GSE214665).

## Author contributions

M. Shiraishi conceived and designed the study. M. Shiraishi conducted most of the experiments and acquired the data with support from A. Yamaguchi and K. Suzuki. All three authors participated in the analysis and interpretation of the data. M. Shiraishi wrote and edited the manuscript with input from A. Yamaguchi and K. Suzuki.

## Acknowledgements

This project was supported by SENSHIN Medical Research Foundation, a Grant-in-Aid for Scientific Research (C), TERUMO LIFE SCIENCE FOUNDATION, Suzuken Memorial Foundation, THE KATO MEMORIAL TRUST FOR NAMBYO RESEARCH, Bristol Myers Squibb Grants for non-clinical. We thank Ms. Wendy Alexander-Adams for her assistance in reporting our findings in English.

---

**Supplementary Fig. 1** Development of our three-part in vitro myocardial fibrosis model.

*a*, Schematic of *in vitro* representations of normal myocardium, fibrotic myocardium following hemodynamic loading (myocardium following HL), and myocardium with reduced fibrosis following hemodynamic loading reduction (myocardium following HLR). To establish the model, we investigated the effect of medium substrate and chemical factors (i.e., TGFβ) on differentiation of the fibroblasts. In substrate with a modulus mimicking ECM of normal myocardium (Low*E*→Low*E*), cells were passaged every 48 hours in at 16 kPa. In substrates mimicking ECM of myocardium following HL (High*E* and/or TGFβ→High*E* and/or TGFβ), cells were passaged every 48 hours at 64 kPa. In substrate mimicking ECM of myocardium following HLR (High*E* and/or TGFβ→Low*E*), cells were incubated in high-elasticity medium at 64 kPa for 48 hours and then in low-elasticity medium at 16 kPa for 48 hours.

*b*, qRT-PCR revealed that change from Low*E* to High*E* conditions and/or addition of TGFβ to successive cultures upregulated expression of *αSMA*, a marker of fibroblast activation, and further upregulation of *αSMA* was observed in passages under the same conditions. To the contrary, *αSMA* expression was significantly reduced in cells with change from a High*E*+TGFβ condition to a Low*E* condition in the second succession.

*c,* Representative images of cardiac fibroblasts stained immunocytochemically for vimentin and αSMA. Expression of αSMA was comparable to that evaluated by qRT-PCR. Scale bars: 50 μm (low magnification), 10 μm (high magnification). *b*, *c*, *n=*4 biologically independent samples per culture condition; mean±s.e.m. values are shown, *b*, \**P*<0.05 versus Low*E* or Low*E*→Low*E* culture condition;

^#^*P*<0.05 versus 1^st^ succession; ^†^*P*<0.05 versus 2^nd^ succession; *t*-test. *c*, Scale bars: 50 μm (low magnification), 10 μm (high magnification), \**P*<0.05 versus Low*E*→Low*E* group; ^#^*P*<0.05 versus fibrotic heart group; *t*-test.

**Supplementary Fig. 2** Bioinformatics analysis of cultured fibroblasts in myocardial fibrosis pathological model.

*a,* Top 20 statistically significant enriched molecules are displayed in the heatmap. The heatmap cells are colored according to their *P* values, white cells indicate the lack of enrichment for that term in the corresponding gene list.

*b,* Heart map shows statistically significant enriched terms.

*c, d,* Representative terms from the full cluster in Set A (*b*) and Set B (*c*) were selected and converted into a network layout. More specifically, each term is represented by a circle node, where its size is proportional to the number of input genes fall into that term, and its color represent its cluster identity (i.e., nodes of the same color belong to the same cluster). Terms with a similarity score >0.3 are linked by an edge. The network is visualized with Cytoscape (v3.1.2) with “force-directed” layout and with edge bundled for clarity. One term from each cluster is selected to have its term description shown as label. Label details in Set A and Set B are listed in the Supplementary Table 1.

Supplementary Fig. 3 Inhibition of the apoptotic signaling pathway affects cardiac fibroblast differentiation in the pathological model of myocardial fibrosis.

*a,* Caspase 3 inhibitor Z-DEVD-FMK or vehicle was added to normal myocardium (Low*E*→Low*E*), fibrotic myocardium following hemodynamic loading (myocardium following HL: High*E*+TGFβ→High*E*+TGFβ), and myocardium with reduced fibrosis following hemodynamic loading reduction (myocardium following HLR: High*E*+TGFβ→Low*E*) model.

*b,* Representative images of cardiac fibroblasts stained immunocytochemically for vimentin and αSMA. Z-DEVD-FMK markedly increased activation of cardiac fibroblasts (ratio of vimentin^+^ and αSMA^+^ myofibroblasts to vimentin^+^ fibroblasts) in myocardium following HL.

*c,* Representative images of cardiac fibroblasts stained immunocytochemically for MMP2 and MMP9. Z-DEVD-FMK markedly attenuated differentiation into the fibrinolytic form of cardiac fibroblasts (ratio of vimentin^+^ and MMP2^+^ or MMP9^+^ myofibroblasts to vimentin^+^ fibroblasts) myocardium following HLR. *b*, *c*, Scale bars: 50 μm (low magnification), 10 μm (high magnification), *n=*6 biologically independent samples per group, Mean±s.e.m. values are shown;

\**P*<0.05 vehicle versus Z-DEVD-FMK; ^#^*P*<0.05 versus Low*E*→Low*E*; ^‡^*P*<0.05 versus Low*E*→Low*E*+Z-DEVD-FMK; ^†^*P*<0.05 versus High*E*+TGFβ→High*E*+TGFβ; ^§^*P*<0.05 versus High*E*+TGFβ→High*E*+TGFβ+Z-DEVD-FMK; one-way ANOVA.

Supplementary Fig. 4 Intrinsic pathway of apoptosis is associated with differentiation of cultured cardiac fibroblasts.

*a,* Schematic representation and overview of the apoptotic signaling pathway.

The intrinsic pathway of apoptosis initiates apoptosis by inducing mitochondrial outer membrane permeability (MOMP). The intrinsic pathway mediates activation of caspase-9 and ultimately leads to cell death via activation of caspase-3 and caspase-7. In the intrinsic pathway of apoptosis, the apoptotic threshold is set by the dynamic interaction of B-cell/CLL lymphoma 2 (BCL-2) family members at the mitochondrial outer membrane^16^. BCL-associated X protein (BAX) and BCL-2 homologous antagonist/killer (BAK) initiate apoptosis via MOMP. The activity of these effectors is regulated by BCL-2-like protein 11 (BCL2L11 [also known as BIM]), p53 upregulated modulator of apoptosis (PUMA) and BH3-interacting domain death agonist (BID)^16^. BCL-2, B-cell lymphoma-extra large (BCL-XL) and myeloid cell leukemia 1 (MCL1) prevent MOMP by directly binding to and inhibiting both effectors and activators. BCL2-associated agonist of cell death (BAD) promotes apoptosis by binding and blocking pro-survival proteins. The extrinsic pathway of apoptosis is activated by the binding of extracellular death ligands such as FAS ligand (FASL), tumor necrosis factor (TNF)^16^. In this pathway, apoptosis-promoting signals are transmitted through the death-inducing signaling complex (DISC). This complex activates caspase-8 and caspase-10, and both caspases initiate apoptosis via activation of procaspase-3^16^.

*b*, qRT-PCR revealed tumor necrosis factor alpha (*TNFα*) expression was extremely low and not significantly different between groups. Fas ligand (*FasL*) expression also did not differ significantly between groups.

*c*, Gene expression of tumor necrosis factor alpha receptor 1 (*TNFR1*), tumor necrosis factor alpha receptor 2 (*TNFR2*), and Fas receptor (*FasR*) in fibrotic myocardium following hemodynamic loading (myocardium following HL: High*E*+TGFβ→High*E*+TGFβ) and myocardium with reduced fibrosis following hemodynamic loading reduction (myocardium following HLR: High*E*+TGFβ→Low*E*) was unchanged or significantly reduced compared to those in normal myocardium (Low*E*→Low*E*).

*d,* Compared to the intimate associations of *Bim* and *Caspase-3* with apoptosis-related gene clusters, the association between *Selenbp1* and apoptosis-related gene clusters has not been revealed yet.

Scrm, scrambled; SBP1, Selenbp1; *b, c, n=*4 biologically independent samples per group, Mean±s.e.m. values are shown; \**P*<0.05 scrm siRNA versus SBP1 siRNA; ^#^*P*<0.05 versus Low*E*→Low*E* (scrm siRNA); ^‡^*P*<0.05 versus Low*E*→Low*E* (SBP1 siRNA); ^†^*P*<0.05 versus High*E*+TGFβ→High*E*+TGFβ (scrm siRNA); ^§^*P*<0.05 versus High*E*+TGFβ→High*E*+TGFβ (SBP1 siRNA); one-way ANOVA.

**Supplementary Fig. 5** In vitro Selenbp1 knockdown influences cardiac fibroblast differentiation in myocardial fibrosis.

*a, Scrm siRNA or SBP1 targeting* siRNA mixed with *in vitro* transfection reagent was added to normal myocardium (Low*E*→Low*E* substrate), myocardium following HL (fibrotic myocardium; High*E*+TGFβ→High*E*+TGFβ substrate), and myocardium following HLR: High*E*+TGFβ→Low*E* substrate).

*b,* Representative images of cardiac fibroblasts stained immunocytochemically for vimentin and αSMA. *SBP1* siRNA transfection markedly increased activation of cardiac fibroblasts (ratio of vimentin^+^ and αSMA^+^ myofibroblasts to vimentin^+^ fibroblasts) in myocardium following HL.

*c,* Representative images of cardiac fibroblasts stained immunocytochemically for MMP2 and MMP9. *SBP1* siRNA transfection markedly attenuated differentiation into the fibrinolytic form of cardiac fibroblasts (ratio of vimentin^+^ and MMP2^+^ or MMP9^+^ myofibroblasts to vimentin^+^ fibroblasts) in myocardium following HLR.

*b*, *c*, *n=*6 biologically independent samples per group and mean±s.e.m. values are shown; *b*, *c*, Scale bars: 50 μm (low magnification), 10 μm (high magnification); \**P*<0.05 scrm siRNA versus *SBP1* siRNA; ^#^*P*<0.05 versus Low*E*→Low*E* (scrm siRNA); ^‡^*P*<0.05 versus Low*E*→Low*E* (*SBP1* siRNA);

^†^*P*<0.05 versus High*E*+TGFβ→High*E*+TGFβ (scrm siRNA); ^§^*P*<0.05 versus

High*E*+TGFβ→High*E*+TGFβ (*SBP1* siRNA); one-way ANOVA.

*Supplementary Fig. 6 Supplementary data on left ventricular volume and myocardial weight*.

*a*, Echocardiography showed decreased LVEDV, LVESV, and SV in TAC-subjected heart and improvement of these variables in Heart Re-TAC. *SBP1* knockdown further decreased LVEDV, LVESV, and SV in TAC-subjected heart but inhibited improvement of these variables in Heart Re-TAC.

*b*, Heart size corrected for body size; ratio of heart weight to body weight (mg/g), and ratio of heart weight to tibial length (mg/mm). TAC enlarged myocardium, and release from TAC reduced the hypertrophy. *SBP1* knockdown contributed to the hypertrophy and prolonged the hypertrophy even after TAC release.

SV, stroke volume; BW, body weight; HW, heart weight; TL, tibia length; *a*, *n*=6 animals per group, and mean±s.e.m. values shown; *b*, *n*=6 animals per group, and mean±s.e.m. values are shown; \**P*<0.05 Scrm siRNA versus *SBP1* siRNA; ^#^*P*<0.05 versus normal heart (scrm siRNA); ^‡^*P*<0.05 versus normal heart (*SBP1* siRNA); ^†^*P*<0.05 versus TAC-subjected heart (Scrm siRNA);

^§^*P*<0.05 versus TAC-subjected heart (*SBP1* siRNA); one-way ANOVA.

**Supplementary Table 1.**
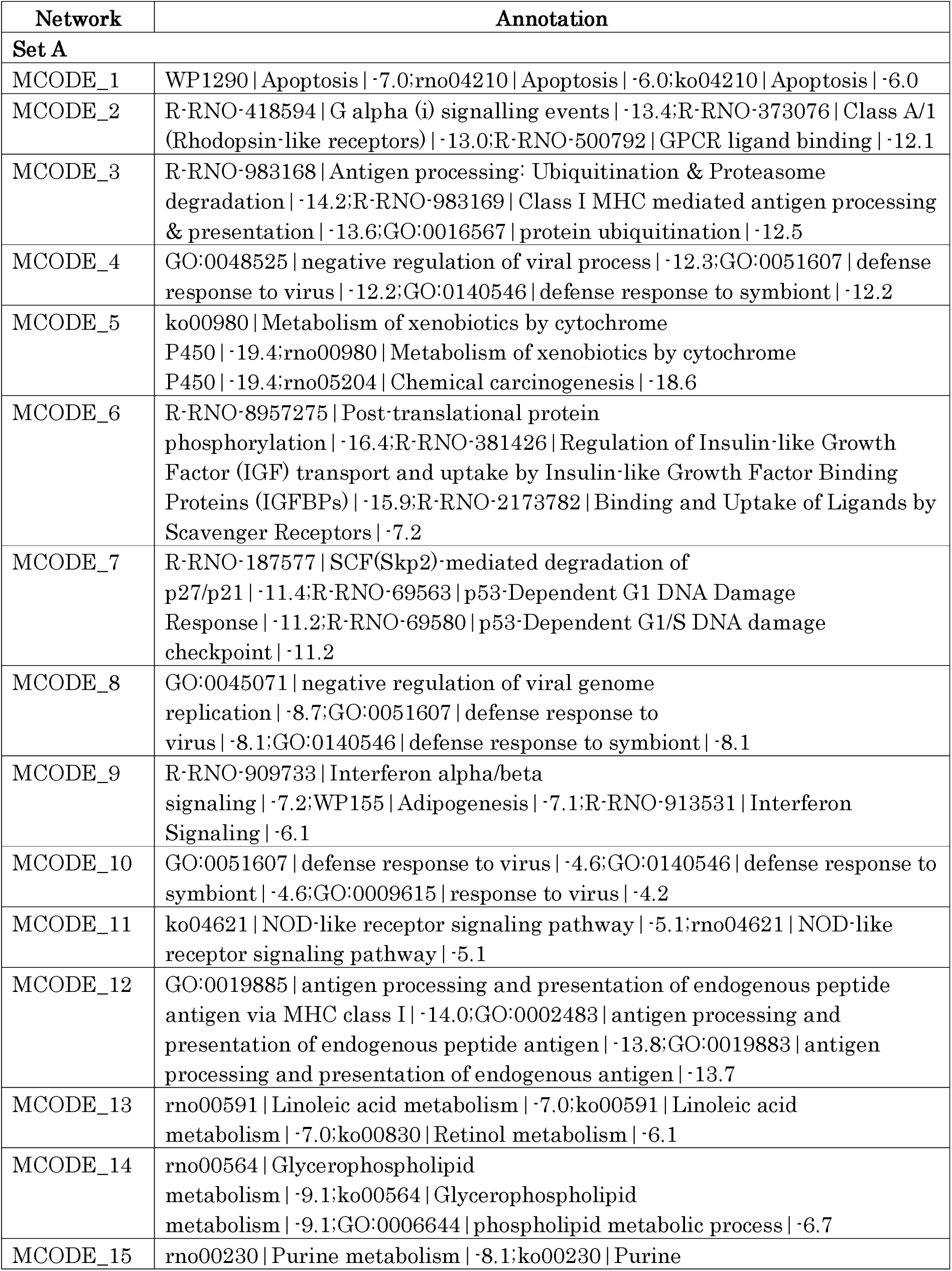
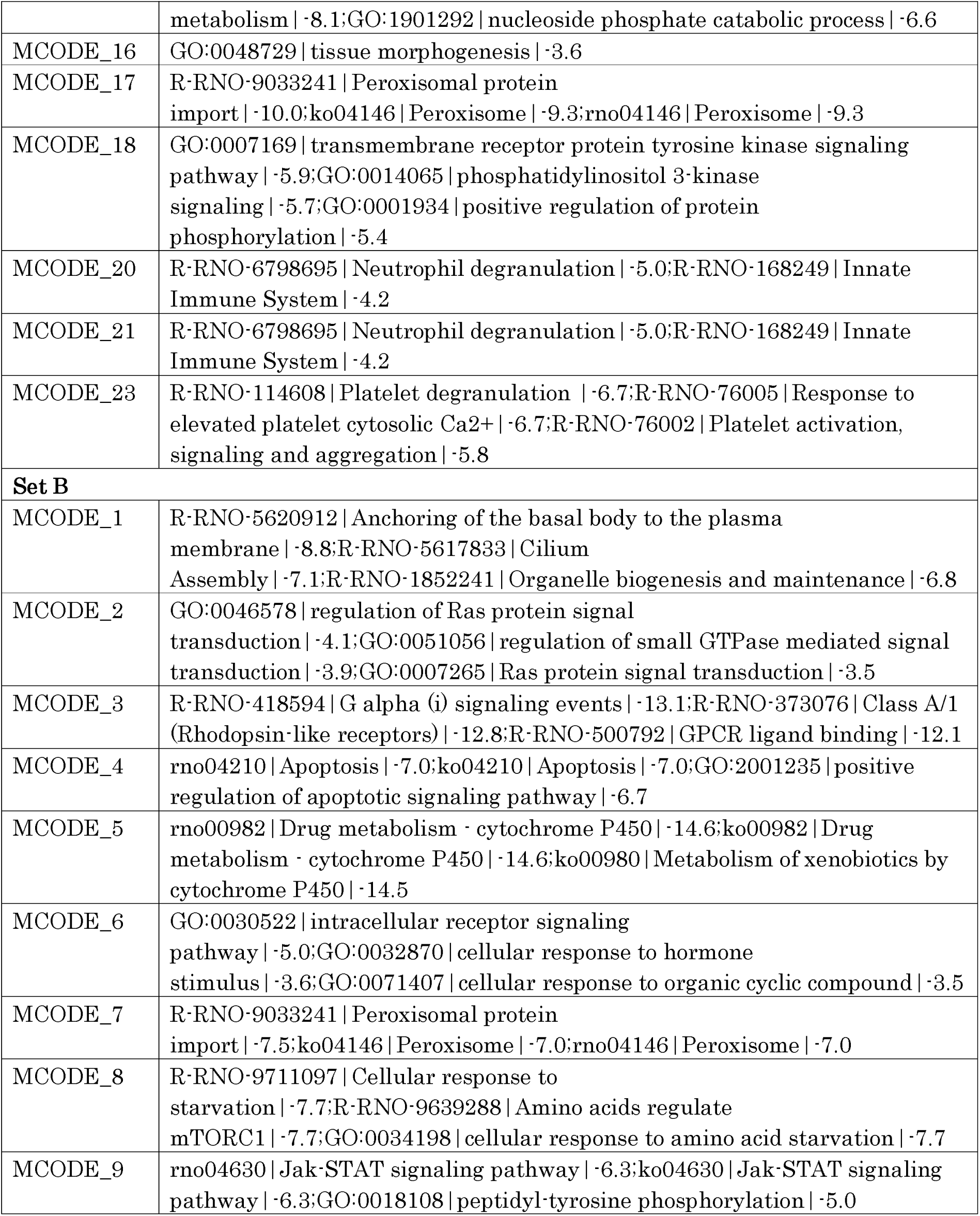
Functional classification of the genes in Set A and Set B.

| Network | Annotation |
| --- | --- |
| **Set A** |  |
| MCODE_1 | WP1290 \| Apoptosis \| ·7.0; rno04210 \| Apoptosis \| ·6.0; ko04210 \| Apoptosis \| ·6.0 |
| MCODE_2 | R-RNO-418594 \| G alpha (i) signalling events \| ·13.4; R-RNO-373076 \| Class A/1 (Rhodopsin-like receptors) \| ·13.0; R-RNO-500792 \| GPCR ligand binding \| ·12.1 |
| MCODE_3 | R-RNO-983168 \| Antigen processing; Ubiquitination & Proteasome degradation \| ·14.2; R-RNO-983169 \| Class I MHC mediated antigen processing & presentation \| ·13.6; GO:0016567 \| protein ubiquitination \| ·12.5 |
| MCODE_4 | GO:0048525 \| negative regulation of viral process \| ·12.3; GO:0051607 \| defense response to virus \| ·12.2; GO:0140546 \| defense response to symbiont \| ·12.2 |
| MCODE_5 | ko00980 \| Metabolism of xenobiotics by cytochrome P450 \| ·19.4; rno00980 \| Metabolism of xenobiotics by cytochrome P450 \| ·19.4; rno05204 \| Chemical carcinogenesis \| ·18.6 |
| MCODE_6 | R-RNO-8957275 \| Post-translational protein phosphorylation \| ·16.4; R-RNO-381426 \| Regulation of Insulin-like Growth Factor (IGF) transport and uptake by Insulin-like Growth Factor Binding Proteins (IGFBPs) \| ·15.9; R-RNO-2173782 \| Binding and Uptake of Ligands by Scavenger Receptors \| ·7.2 |
| MCODE_7 | R-RNO-187577 \| SCF(Skp2)-mediated degradation of p27/p21 \| ·11.4; R-RNO-69563 \| p53-Dependent G1 DNA Damage Response \| ·11.2; R-RNO-69580 \| p53-Dependent G1/S DNA damage checkpoint \| ·11.2 |
| MCODE_8 | GO:0045071 \| negative regulation of viral genome replication \| ·8.7; GO:0051607 \| defense response to virus \| ·8.1; GO:0140546 \| defense response to symbiont \| ·8.1 |
| MCODE_9 | R-RNO-909733 \| Interferon alpha/beta signaling \| ·7.2; WP155 \| Adipogenesis \| ·7.1; R-RNO-913531 \| Interferon Signaling \| ·6.1 |
| MCODE_10 | GO:0051607 \| defense response to virus \| ·4.6; GO:0140546 \| defense response to symbiont \| ·4.6; GO:0009615 \| response to virus \| ·4.2 |
| MCODE_11 | ko04621 \| NOD-like receptor signaling pathway \| ·5.1; rno04621 \| NOD-like receptor signaling pathway \| ·5.1 |
| MCODE_12 | GO:0019885 \| antigen processing and presentation of endogenous peptide antigen via MHC class I \| ·14.0; GO:0002483 \| antigen processing and presentation of endogenous peptide antigen \| ·13.8; GO:0019883 \| antigen processing and presentation of endogenous antigen \| ·13.7 |
| MCODE_13 | rno00591 \| Linoleic acid metabolism \| ·7.0; ko00591 \| Linoleic acid metabolism \| ·7.0; ko00830 \| Retinol metabolism \| ·6.1 |
| MCODE_14 | rno00564 \| Glycerophospholipid metabolism \| ·9.1; ko00564 \| Glycerophospholipid metabolism \| ·9.1; GO:0006644 \| phospholipid metabolic process \| ·6.7 |
| MCODE_15 | rno00230 \| Purine metabolism \| ·8.1; ko00230 \| Purine |
|  | metabolism \| ·8.1;GO:1901292 \| nucleoside phosphate catabolic process \| ·6.6 |
| MCODE_16 | GO:0048729 \| tissue morphogenesis \| ·3.6 |
| MCODE_17 | R·RNO-9033241 \| Peroxisomal protein import \| ·10.0;ko04146 \| Peroxisome \| ·9.3;rno04146 \| Peroxisome \| ·9.3 |
| MCODE_18 | GO:0007169 \| transmembrane receptor protein tyrosine kinase signaling pathway \| ·5.9;GO:0014065 \| phosphatidylinositol 3-kinase signaling \| ·5.7;GO:0001934 \| positive regulation of protein phosphorylation \| ·5.4 |
| MCODE_20 | R·RNO-6798695 \| Neutrophil degranulation \| ·5.0;R·RNO-168249 \| Innate Immune System \| ·4.2 |
| MCODE_21 | R·RNO-6798695 \| Neutrophil degranulation \| ·5.0;R·RNO-168249 \| Innate Immune System \| ·4.2 |
| MCODE_23 | R·RNO-114608 \| Platelet degranulation \| ·6.7;R·RNO-76005 \| Response to elevated platelet cytosolic Ca<sup>2+</sup> \| ·6.7;R·RNO-76002 \| Platelet activation, signaling and aggregation \| ·5.8 |
| **Set B** |  |
| MCODE_1 | R·RNO-5620912 \| Anchoring of the basal body to the plasma membrane \| ·8.8;R·RNO-5617833 \| Cilium Assembly \| ·7.1;R·RNO-1852241 \| Organelle biogenesis and maintenance \| ·6.8 |
| MCODE_2 | GO:0046578 \| regulation of Ras protein signal transduction \| ·4.1;GO:0051056 \| regulation of small GTPase mediated signal transduction \| ·3.9;GO:0007265 \| Ras protein signal transduction \| ·3.5 |
| MCODE_3 | R·RNO-418594 \| G alpha (i) signaling events \| ·13.1;R·RNO-373076 \| Class A/1 (Rhodopsin-like receptors) \| ·12.8;R·RNO-500792 \| GPCR ligand binding \| ·12.1 |
| MCODE_4 | rno04210 \| Apoptosis \| ·7.0;ko04210 \| Apoptosis \| ·7.0;GO:2001235 \| positive regulation of apoptotic signaling pathway \| ·6.7 |
| MCODE_5 | rno00982 \| Drug metabolism · cytochrome P450 \| ·14.6;ko00982 \| Drug metabolism · cytochrome P450 \| ·14.6;ko00980 \| Metabolism of xenobiotics by cytochrome P450 \| ·14.5 |
| MCODE_6 | GO:0030522 \| intracellular receptor signaling pathway \| ·5.0;GO:0032870 \| cellular response to hormone stimulus \| ·3.6;GO:0071407 \| cellular response to organic cyclic compound \| ·3.5 |
| MCODE_7 | R·RNO-9033241 \| Peroxisomal protein import \| ·7.5;ko04146 \| Peroxisome \| ·7.0;rno04146 \| Peroxisome \| ·7.0 |
| MCODE_8 | R·RNO-9711097 \| Cellular response to starvation \| ·7.7;R·RNO-9639288 \| Amino acids regulate mTORC1 \| ·7.7;GO:0034198 \| cellular response to amino acid starvation \| ·7.7 |
| MCODE_9 | rno04630 \| Jak-STAT signaling pathway \| ·6.3;ko04630 \| Jak-STAT signaling pathway \| ·6.3;GO:0018108 \| peptidyl-tyrosine phosphorylation \| ·5.0 |

